# Region-Specific PSD-95 Interactomes Contribute to Functional Diversity of Excitatory Synapses in Human Brain

**DOI:** 10.1101/2020.05.04.076844

**Authors:** Adam J. Funk, Guillaume Labilloy, James Reigle, Rawan Alnafisah, Michael R. Heaven, Rosalinda C. Roberts, Behrouz Shamsaei, Kenneth D. Greis, Jaroslaw Meller, Robert E. McCullumsmith

## Abstract

The overarching goal of this exploratory study is to link subcellular microdomain specific protein-protein interactomes with big data analytics. We isolated postsynaptic density-95 (PSD-95) complexes from four human brain regions and compared their protein interactomes using multiple bioinformatics techniques. We demonstrate that human brain regions have unique postsynaptic protein signatures that may be used to interrogate perturbagen databases. Assessment of our hippocampal signature using the iLINCS database yielded several compounds with recently characterized “off target” effects on protein-protein interactions in the posynaptic density compartment.

## Introduction

The human brain is comprised of billions of neurons that form networks of connections within and between brain regions [1]. These connections facilitate neuroplastic events that underlie learning and memory, critical aspects of cognitive function often perturbed in neuropsychiatric illnesses [2, 3]. Neuronal signaling is mediated by fast and slow transmission events, encompassing receptors, ligands, ions, enzymes, and other substrates [1, 4-7]. These elements are spatially arranged in subcellular microdomains, facilitating juxtaposition of proteins that coordinate various biological processes. For example, synaptic transmission is modulated via release of neurotransmitter into the synaptic cleft, where receptors are activated and the postsynaptic cell modulated via electrical and chemical signals [1, 4-6].

The pre- and postsynaptic compartments include highly specialized protein clusters, with elegant and complex regulatory mechanisms that traffick proteins to and from these zones. In particular, postsynaptic densities are microdomains comprised of about 1000 unique proteins that are interacting with one another via specialized scaffolding molecules [8]. Postsynaptic density-95 (PSD-95) is a multipotent scaffolding, trafficking, and clustering protein which links glutamate receptors, signaling molecules, and other structural proteins at postsynaptic sites. More than 95% of PSD-95 expression is localized to excitatory synapses, and it is the most abundant scaffolding protein within the postsynaptic density [9-11].

Highly linked proteins, or hubs, are usually protected from null mutations due to the overrepresentation of deleterious functional outcome [12]. PSD-95 is a highly interconnected hub protein with known functional consequences if manipulated. Altering the expression of PSD-95 protein significantly changes synaptic structure and function, including molecular correlates of learning and memory, long-term potentiation (LTP) and long-term depression (LTD). For example, PSD-95 overexpression increases the amplitude of excitatory postsynaptic currents (EPSCs), to the extent that additional stimulus is unable to further strengthen LTP [13-17]. In contrast, PSD-95 knockdown disrupts postsynaptic density structure and suppresses EPSCs, consistent with a role for PSD-95 in synaptic fidelity [18-20]. Further highlighting the importance of PSD-95 in the synapse, knockout of PSD-95 in mice and genomic variants within postsynaptic density hub proteins are particularly implicated in neuropsychiatric disorders such as schizophrenia and autism [21, 22].

To investigate excitatory postsynaptic protein hubs, we targeted PSD-95 due to its high level of localization to the postsynaptic density (∼95%), its well-characterized role as a hub protein, and the large number of constituents in its protein interactome (∼1000) [23-25]. We designed a series of experiments to examine brain region-specific excitatory postsynaptic proteomes to answer the following questions. 1) Do postsynaptic protein-protein interactions differ by brain region? 2) Can we accurately detect and separate samples based on these differences? and 3) Can region-specific protein interactomes inform functional differences? For these experiments we used four human brain regions (anterior cingulate cortex (ACC), dorsolateral prefrontal cortex (DLPFC), hippocampus (HPC), and superior temporal gyrus (STG)) from 3 well-matched control male subjects from the Alabama Brain Collection. A total of 289 proteins were identified as part of a PSD-95 protein interactome that were present across all four brain regions. We hypothesize that there is region-specific heterogeneity in the composition of excitatory postsynaptic microdomains, reflecting the differences in neuroplastic function between human brain regions.

### Experimental Procedures

#### Workflow overview

Human brain tissue from three male subjects from four brain regions was processed for affinity purification of PSD-95 protein complexes (Table 1). We confirmed PSD-95 capture and enrichment from each sample using Western blot analyses. For quality control, individual biological samples were pooled together based on region, PSD-95 complexes were affinity purified, processed for mass spectrometry, and run in technical triplicate. From the pooled PSD-95 affinity purified complexes we identified more than 500 unique peptides in each pooled sample from each region using data-independent acquisition, as well as shotgun proteomic analyses, confirming robust PSD-95 complex capture (data not shown). Next, individual samples (n = 3 biological replicates for each of four brain regions) were run through our PSD-95 affinity and mass spectrometry protocol in triplicate (3 technical replicates x 12 biological samples for 36 runs total). We then subtracted any non-specifically captured peptides identified by our IgG control studies that were performed in parallel to PSD-95 affinity purification. To account for processing and loading variability, data were normalized within each of the 36 runs to the most abundant PSD-95 peptide (IIPGGAAAQDGR) across all samples. Only peptides that were present in at least 2 of 3 technical replicates were carried forward (peptides missing a technical replicate were replaced by imputation), then all 36 runs were quantile normalized [26, 27]. We used multiple clustering approaches to identify peptides which contributed to brain region specificity, unsupervised clustering (Pearson correlation) and a weakly supervised clustering protocol (non-negative matrix factorization, NMF), using the same peptides for both clustering analyses [28-30]. T-statistic and coefficient values from unsupervised and NMF clustering were generated for each peptide and a consensus peptide profile, or signature, was generated based on common input peptides across all 4 regions. The region-specific concensus signatures were subjected to traditional and advanced bioinformatics analyses to identify pathways, processes, and compounds associated with the signatures for each brain region.

**Table 1.** PSD-95 interactome subject demographics

| Subject | 1 | 2 | 3 |
| --- | --- | --- | --- |
| Dx | Normal | Normal | Normal |
| Age | 73 | 71 | 71 |
| pH | 6.4 | 6.4 | 6.4 |
| PMI hrs | 17 | 13 | 20 |
| Sex | Male | Male | Male |
| Ethnicity | Caucasian | Caucasian | Caucasian |

#### Tissue preparation

Deidentified cortical tissue samples from 3 matched male subjects were obtained from the Alabama Brain Collection (ABC). The ABC obtained informed consent from survivors of all donors, as well as IRB approval (at the University of Alabama Birmingham) for human brain collection. Broadmann areas 46, 32, 22, and hippocampus were dissected from coronal slabs and stored at 80C. For each region, frozen tissue was thawed at −20°C for 1 hour then sectioned onto glass slides at 50um. Ten 50 um sections were used for each region. Grey matter was harvested from each slide, combined by region, and homogenized 30x on ice with a Teflon coated dounce homogenizer in 1ml of 5mM Tris HCl, 0.32 M Sucrose, with 1% Halt protease and phosphatase inhibitor, pH 7.4. Protein concentration was measured with a Pierce BCA kit. All samples were normalized for protein concentration. TritonX-100 and SDS were then added to the homogenate at a final concentration of 1% v/v and 0.5% v/v, respectively, and then rotated overnight at 4°C. Lysates were immediately used for affinity purification and remaining sample was stored at −80°C.

#### Affinity purification

We used a mouse anti-PSD-95 antibody (Millipore, catalogue # MAB1596) to capture PSD-95 protein complexes from individual samples. We verified the specificity of this antibody using mass spectrometry analysis of PSD-95 peptides captured by affinity purification vs isotype IgG control (Figure S1). Our LCMS protocol yielded ∼65% coverage of the PSD-95 protein (Figure S1C). 5ug of PSD-95 antibody was coupled per 1mg of Dynabeads (Life Technologies) according to the antibody coupling kit protocol (#14311D). For each sample, 1000ul of 10mg/ml antibody coupled beads were washed 2x 1ml with ice cold 1x PBST (#9809S, Cell Signaling) then incubated with 650ug of human brain lysate brought to a final volume of 1000ul with ice cold 1x PBST for 1 hour at room temperature. The supernatant was removed and saved for western blot analysis and the beads were washed 4 x 10 minutes at room temperature in 1ml ice cold 1x PBST. Captured protein complexes were eluted with 30ul of 1N Ammonium Hydroxide (#320145, Sigma), 5mM EDTA, pH 12 for 10 minutes at room temperature. 6ul of 6x protein denaturing buffer (4.5% SDS, 15% β-mercaptoethanol, 0.018% bromophenol blue, and 36% glycerol in 170 mM Tris-HCl, pH 6.8) was added to each sample elution. The eluted samples were heated at 70°C for 10 minutes then processed for mass spectrometry.

#### Antibody control studies

We performed parallel studies using species matched preimmune IgG. All apsects of the affinity purification described above were duplicated as described above. Three affinity purified IgG control samples (biological replicates) from each region were pooled. We ran three technical replicates of one pooled sample for each brain region through the mass spectrometry protocol. Peptides detected in at least 2 out of 3 technical replicates were designated as non-specific, and subtracted from our PSD95 affinity purification mass spectrometry dataset analyses.

#### Mass Spectrometry sample preparation

All samples were loaded on a 1.5 mm, 4-12% Bis-Tris Invitrogen NuPage gel (NP0335BOX) and electrophoresed in 1x MES buffer (NP0002) for 10 minutes at 180v. The gel was fixed in 50% ethanol/10% acetic acid overnight at RT, then washed in 30% ethanol for 10 min followed by two 10 min washes in MilliQ water (MilliQ Gradient system). The lanes were harvested, cut into small (∼2mm) squares, and subjected to in-gel tryptic digestion and peptide recovery, as described previously [31]. Samples were resuspended in 0.1% formic acid.

#### Nano liquid chromatography coupled electrospray tandem mass spectrometry (nLC-ESI-MS/MS)

nLC-ESI-MS/MS analyses were performed on a 5600+ QTOF mass spectrometer (Sciex, Toronto, On, Canada) interfaced to an Eksigent (Dublin, CA) nanoLC ultra nanoflow system. Samples were run consecutively then repeated for technical replicates 2 and 3, as previously described [31]. Breifly, peptides were loaded onto an IntegraFrit Trap Column at 2 µl/min in formic acid/H2O 0.1/99.9 (v/v) for 15 min to desalt and concentrate the samples then the trap-column was switched to align with the analytical column. Peptides were eluted using a variable mobile phase (MP) gradient from 95% phase A (Formic acid/H2O 0.1/99.9, v/v) to 40% phase B (Formic Acid/Acetonitrile 0.1/99.9, v/v) for 70 min, from 40% phase B to 85% phase B for 5 min and then keeping the same mobile phase composition for 5 additional min at 300 nL/min. The nLC effluent was ionized and sprayed into the mass spectrometer using NANOSpray® III Source (Sciex). Ion source gas 1 (GS1), ion source gas 2 (GS2) and curtain gas (CUR) were respectively kept at 8, 0 and 35 vendor specified arbitrary units.

The mass spectrometer method was operated in positive ion mode and the interface heater temperature and ion spray voltage were kept at 150°C, and at 2.6 kV, respectively. The data was recorded using Analyst-TF (version 1.7) software.

#### Data independent acquisition (DIA) and shotgun (information dependent acquisition, IDA) methods

The DIA method was set to go through 1757 cycles for 99 minutes, where each cycle performed one TOF-MS scan type (0.25 sec accumulation time, in a 550.0 to 830.0 m/z window) followed by 56 sequential overlapping windows of 6 Daltons each. Note that the Analyst software automatically adds 1 Dalton to each window to provide overlap, thus an input of 5 Da in the method set up window results in an overlapping 6 Da collection window width (e.g. 550-556, then 555-561, 560-566, etc). Within each window, a charge state of +2, high sensitivity mode, and rolling collision energy with a collision energy spread (CES) of 15 V was selected. For shotgun acquisition the mass spectrometer was set to perform one TOF-MS scan type (0.25 sec accumulation time, in a 350 to 1600 m/z window) followed by 50 information dependent acquisition (IDA)-mode MS/MS-scans on the most intense candidate ions having a minimum intensity of 150 counts. Each MS/MS scan was operated under vender specified high-sensitivity mode with an accumulation time of 0.05 secs and a mass tolerance of 100 ppm. Precursor ions selected for MS/MS scans were excluded for 30 secs to reduce the occurrence of redundant peptide sequencing.

#### DIA and IDA data analysis parameters

Proteowizard was used to generate peaks from the raw files using Sciex standard settings [32-34]. Protalizer DIA software (<u>Vulcan Analytical</u>, Birmingham, AL), a spectral library-free DIA analysis package, was used to analyze every DIA file with settings previously described for Sciex 5600+ QTOFs [35, 36]. The Swiss-Prot *Homo Sapien* database, downloaded March 17^th^ 2015, was used as the reference database for all MS/MS searches. A maximum of 2 missed cleavages was allowed for searching based on trypsin cleavage. A precursor and fragment-ion tolerance for QTOF instrumentation was used for the Protalizer Caterpillar spectral-library free identification algorithm as previously described [36, 37]. Peptide retention time and drift adjustment were corrected as previously described [36, 37]. Potential modifications included in the searches were phosphorylation at S, T, and Y residues, N-terminal acetylation, N-terminal loss of ammonia at C residues, and pyroglutamic acid at N-terminal E and Q residues. Carbamidomethylation of C residues was searched as a fixed modification. The maximum valid protein and peptide expectation score from the X! Tandem Sledgehammer search engine (version 2013.09.01.1) used for peptide and protein identification on reconstructed spectra was set to 0.005. The forward and reverse *H. Sapien* Swiss-Prot database was used for identification.

For DIA quantification, the maximum number of b and y series fragment-ion transitions were set to nine excluding those with *m/z* values below 300 and not containing at least 10% of the relative intensity of the strongest fragment-ion assigned to a peptide. A minimum of five fragment-ions were required for a peptide to be quantified. In datasets where a minimum of seven consistent fragment-ions were not detected for the same peptide ion in each of the three files compared in a triplicate analysis, the algorithm identified the file with the largest sum fragment-ion AUC and extracted up to seven of these in the other files using normalized retention time coordinates based on peptides detected by the Caterpillar algorithm in all the files in a dataset. The calculated maximum FDR rate at the peptide and protein level was 1.45% and 9.3% respectively.

#### Data Availability

The raw and processed files have been downloaded to PeptideAtlas and are publicly available using the following link. Chromatograms of all peptides identified are included as web viewable files. http://www.peptideatlas.org/PASS/PASS01361

#### Western blot analyses

A 2ul aliquot was taken from the previously reduced and denatured samples destined for mass spectrometry and processed by SDS-PAGE using Invitrogen (Carlsbad, CA) 1.5mm thick, 4 − 12% gradient gels and transferred to PVDF membranes via BioRad semi-dry transblotters (Hercules, CA). The membranes were blocked with LiCor blocking buffer (Lincoln, NE) for 1 hour at room temperature, then probed for 1 hour at room temperature with primary antisera for PSD-95 (1:1000, Cell Signaling, catalogue # D27E11), Synaptophysin (1:1000, Cell Signaling, catalogue # D40c4), or CaMKIIalpha (1:1000, Santa Cruz Biotechnology, catalogue # sc-376828) diluted in 0.1% Tween LiCor blocking buffer. The membranes were washed twice for 10 minutes each in 1x 0.1% Tween phosphate buffer solution (PBST) then probed with goat anti-mouse or goat anti-rabbit IR-Dye 670 or 800cw-labeled secondary antisera (LiCor) in LiCor blocking buffer (plus 0.1% Tween, 0.01% sodium dodecyl sulfate (SDS)) for 1 hour at room temperature. Washes were repeated after secondary labeling, washing twice for 10 minutes in 1x PBST, and then placed in water. Membranes were imaged using a LiCor Odyssey scanner.

#### Experimental design and statistical rationale

We used area under the curve to quantify the abundance for 424 peptides that were captured as interacting with PSD-95 in four brain-regions in three healthy patients. Each biological sample was run in technical triplicate, resulting in thirty-six measurements that were processed and analyzed using an in house computational pipeline developed using R version 3.1.3. Since the experimental design was optimized for measuring relative, rather than absolute abundance levels, each measurement vector (sample) was first normalized to express peptide abundance levels in relative terms. This is achieved by dividing each peptide’s abundance estimates by the intensity of the PSD-95 IIPGGAAAQDGR peptide. We selected this peptide for its consistency and higher than average intensity across all samples compared to other PSD-95 peptides (data not shown). Note that highly abundant fragments are expected to be quantified more reliably, and therefore higher intensities are more likely to represent the real abundance of a component when compared between samples. Data were then Log2-transformed and, to improve analysis reliability, only peptides with at least two out of three data points per biological sample available for each peptide were considered. In the current work, we focused on PSD-95-associated proteins with a higher abundance in specific regions of the brain, but at same time observed (at lower levels) in multiple regions. Thus, proteins that are not observed in some regions at all (or have too many missing values) were excluded to avoid confounding the results by including false negatives. The rates of false negatives in MS-based proteomics, including those peptide moieties that are not detected because their intensities are too low (given the noise levels), and can be very significant [38]. Therefore, a larger sample size (than used in the present work) would be required to confidently separate biological and technical negatives, which is the subject of future work. Each sample was then quantile-normalized to account for signal amplitude variation across measurements. Remaining missing values were imputed by the mean of the two other data points within three technical replicates per biological sample.

#### Clustering analyses and generation of signatures

Unsupervised clustering was first used in order to identify potential strata of samples with distinct protein expression patterns. Coefficient of variation of each peptide’s abundance levels was used as a measure of variability to rank and select the 200 most variable peptides across all 36 runs. For the purpose of visualization and further analysis, each peptide row was then z-transformed in order to align the mean and amplitude of peptides across samples. Patterns in protein expression data were identified by clustering both peptides and samples with the Pearson correlation based distance function, and visualized using the *Heatplot* function from the *made4* R-package. For further validation and assessment of the effects of potential outliers, Spearman rank correlation was used as an alternative, yielding very similar results (data not shown).

Secondly, NMF decomposition of the fully-normalized matrix was obtained for decomposition dimensions k ranging from 4 to 7 using the “ls-nmf” algorithm after shifting the matrix values by 10 in order to obtain a non-negative input matrix [30]. The decomposition on k=6 vectors appeared to best segregate samples and was therefore selected for further analysis.

Signatures were derived from the two distinct clustering analyses. From the unsupervised cluster, t-tests were performed for each peptide for each group against all the other groups. The 50 most specific peptides for each group defined its signature. On the other hand, the NMF decomposition presented specific relations between HPC and ACC and vector 3 and 1 of the decomposition. Since the relations between the other vectors (2, 4, 5 and 6) with brain regions were less obvious, we focused on the NMF signature for HPC and ACC. We then ranked the peptides (positive) decomposition coefficient and selected the 50 most specific peptides for HPC and ACC. A consensus signature containing overlapping hits from each clustering approach was used for subsequent bioinformatics analyses.

#### Traditional bioinformatics analyses

We first used ToppCluster (https://toppcluster.cchmc.org/) in order to perform gene set enrichment analysis and map pathways associated with region specific PSD-95 interactome signatures [39]. Additional analyses were performed using piNET (http://pinet-server.org/), a tool developed to provide further assessment of region-specific protein networks, including targeted analyses of kinase signaling pathways [40].

#### Advanced bioinformatics analyses

We also used the Library of Integrated Network-based Cellular Signatures (LINCS) (http://ilincs.org) to analyze the connectivity of region specific PSD-95 protein interactome patterns [41-47]. With the goal of identifying connected gene knockdown and small drug-like molecule signatures, we used iLINCS to first retrieve transcriptional signatures of gene knockdowns for genes contributing to HPC versus ACC specific patterns. We then used iLINCS to identify small molecules that result in signatures that are significantly discordant (or reversing these gene knockdown signatures) or concordant with these LINCS knockdown signatures. Only genes that had gene knockdowns available were included (from the top 50 proteins in the overexpression/consensus signature). Vertebral-Cancer of the Prostate (VCAP) cell lines are used as the primary source of knockdowns (if not available, the A375 cell line was used since these two cell lines have the best coverage). Only significant correlations with drug signatures were considered (top 1%). Discordant signatures were primarily from VCAP, with additional cell lines considered as supporting evidence. We then performed literature review of small molecules suggested by our analyses to determine if there is evidence for a mechanism linking our proteomic studies with drugs or perturbagens that yield discordant signatures.

## Results

We developed a method for capturing the PSD-95 protein complexes from human brain by adapting and modifying an approach used in rodent brain (Figure 1)[48]. Using homogenized human frontal cortex tissues, we partially solubilized membrane associated protein complexes by adding sodium dodecyl sulfate (SDS) at a final concentration of 0.5% (Figure 1A). This sufficiently released PSD-95 complexes from membranes and increased enrichment of PSD-95. Increasing concentrations of SDS resulted in reduced capture of PSD-95 (Figure 1B). Notably, representative protein-protein interactions with CaMKIIα remain well preserved using 0.5% SDS, while presynaptic markers (synaptophysin) are not captured. We further characterized PSD-95 affinity purified complexes by coomassie staining of total protein. We show a robust enrichment of PSD-95 protein and its interacting partners, with (apparently) hundreds of abundantly co-purified proteins (Figure 1C). Isotype specific IgG negative control studies demonstrate the specificity of our protocol (Figure 1D).

**Figure 1.**
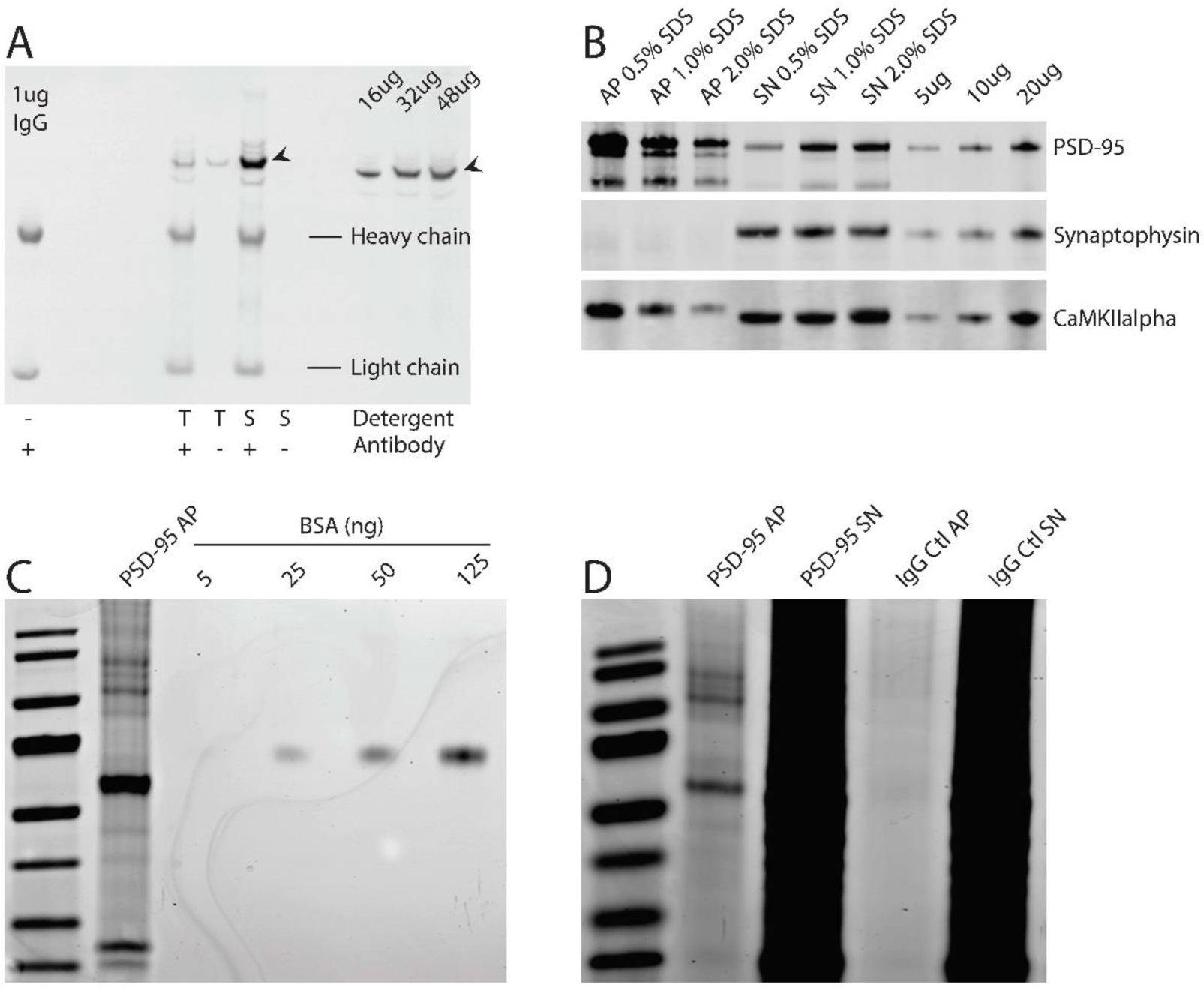
Western blot analysis of PSD-95 affinity purification (AP) using mouse anti-PSD-95 antibody conjugated to magnetic capture beads. **Panel A**: We assessed several detergents, including triton X-100 (T), and sodium dodecyl sulfate (S or SDS), and found that SDS yielded efficient capture of PSD-95, with little or no nonspecific binding in the control lane without antibody (beads alone). The three lanes on the right side of panel A are from whole dorsolateral prefrontal cortex homogenate used as a positive control. **Panel B**: We next determined the optimal concentration of SDS for PSD-95 capture and preservation of the well-characterized PSD-95 interacting protein calcium/calmodulin kinase II alpha (CaMKIIalpha). The presynaptic marker synaptophysin was only present in the PSD-95 AP supernatant (SN). **Panel C**: Coomassie blue staining of SDS-PAGE gels loaded with captured PSD-95 AP sample, bovine serum albumin (BSA), or the PSD-95 AP supernatant. 50ug of mouse anti-PSD-95 antibody conjugated to 10mg of Dynabeads was incubated with 1.4 mg of normal human frontal cortex homogenate for 1 hour at room temperature. The PSD-95 AP yielded discrete, densely stained bands. **Panel D**: Coomassie blue staining of PSD-95 or IgG AP, and the corresponding supernatant for each sample, showing little non-specific binding to the bead-antibody complex.

We next performed PSD-95 affinity purification from 4 brain regions including the anterior cingulate cortex (ACC), dorsolateral prefrontal cortex (DLPFC), hippocampus (HPC), and superior temporal gyrus (STG) from three age-matched elderly human male subjects (Table 1 and Figure 2A and 2B). Samples were processed for mass spectrometry and each biological sample was run three times through an ABSciex 5600+ mass spectrometer in DIA mode (550-830 m/z range with 6m/z windows). Representative chromatograms for 2 abundant PSD-95 peptides (IIPGGAAAQDGR and NTYDVVYLK) are shown to indicate how area under the curve abundance values were generated for each peptide (Figure 2C). The robust reproducibility of our data capture method is represented by the low coefficient of variation displayed by the majority of peptides measured from the technical replicates (Figure 2D).

**Figure 2.**
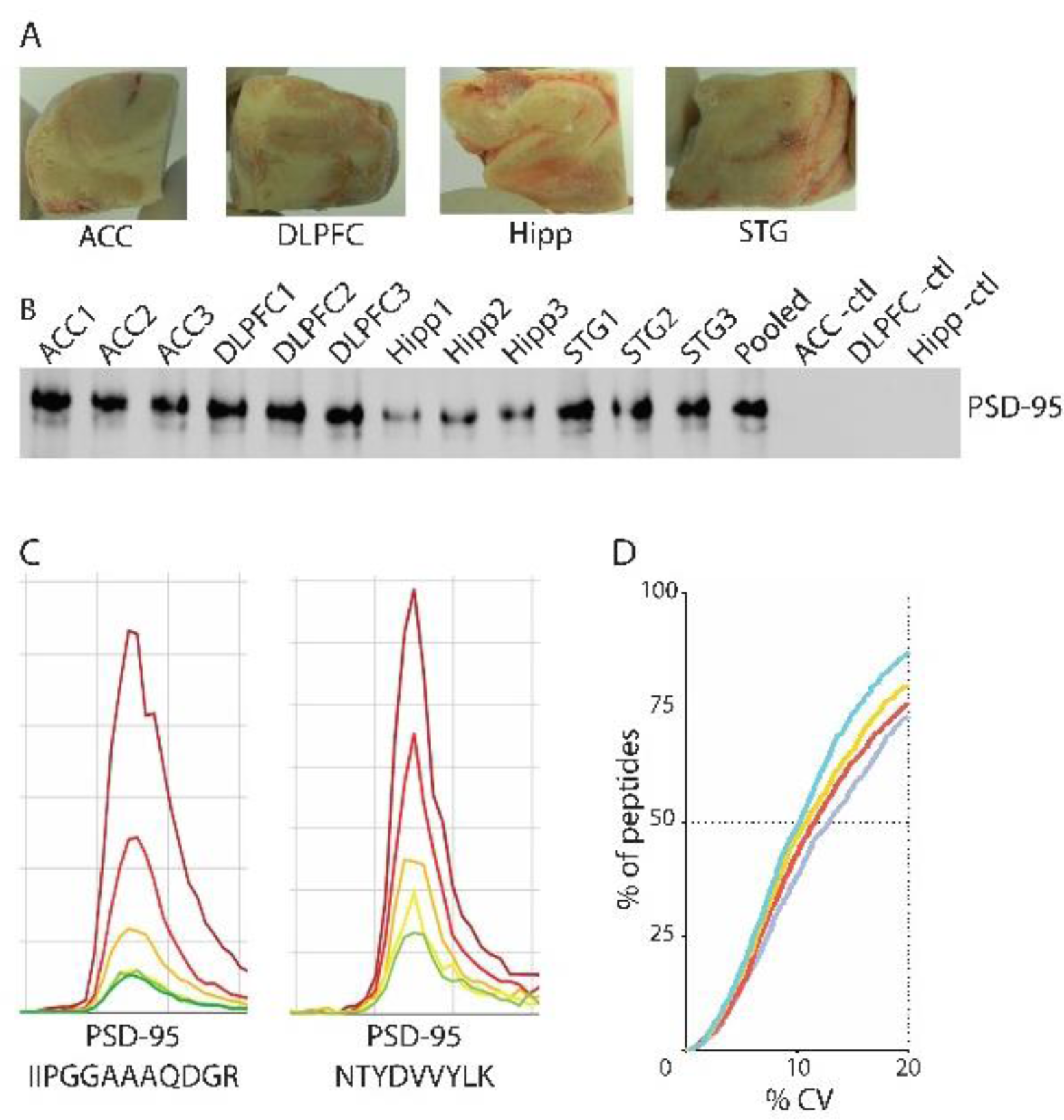
Regional PSD-95 affinity purification of human brain homogenate. **Panel A**: PSD-95 protein interactome was affinity purified (AP) from 4 brain regions, anterior cingulate cortex (ACC), dorsolateral prefrontal cortex (DLPFC), hippocampus (HPC), and superior temporal gyrus (STG) from three normal human male brains. **Panel B**: 2ul of the PSD-95 AP elution for each sample (unpooled) was assessed for PSD-95 protein by western blot analysis. **Panel C**: Data independent acquisition (DIA) mass spectrometry analysis of PSD-95 AP samples yielded hundreds of peptides, including PSD-95. Chromatograms were generated (e.g. PSD-95 peptides, IIPGGAAAQDGR and NTYDVVYLK) and area under the curve was measured for all peptides identified. **Panel D**: Coefficient of variation (% CV) analysis was performed for all peptides with 3 technical replicates from each biological sample (n = 4 regions x 3 subjects = 12 samples). The %CV values from all 3 biological samples were combined into 1 curve for each region.

Peptides detected by LCMS were normalized to the abundance of IIPGGAAAQDGR, then quantile normalized for inter-sample correlation analyses and careful missing-value handling [26, 49-51]. After normalization, we performed unsupervised clustering of all biological and technical replicates using the 200 most variable peptides to determine if normalized peptide abundance signatures could differentiate inter-subject and inter-region differences (Figure 3). Technical replicates clustered together 100% of the time while regional samples showed some degree of overlap (1 DLPFC sample was more closely associated with all 3 ACC rather than the other 2 DLPFC samples, while 1 STG and 1 HPC sample were more closely associated with two DLPFC samples (Figure 3). Of the four regions, the HPC and ACC clustered most consistently using this standard differential expression approach.

**Figure 3.**
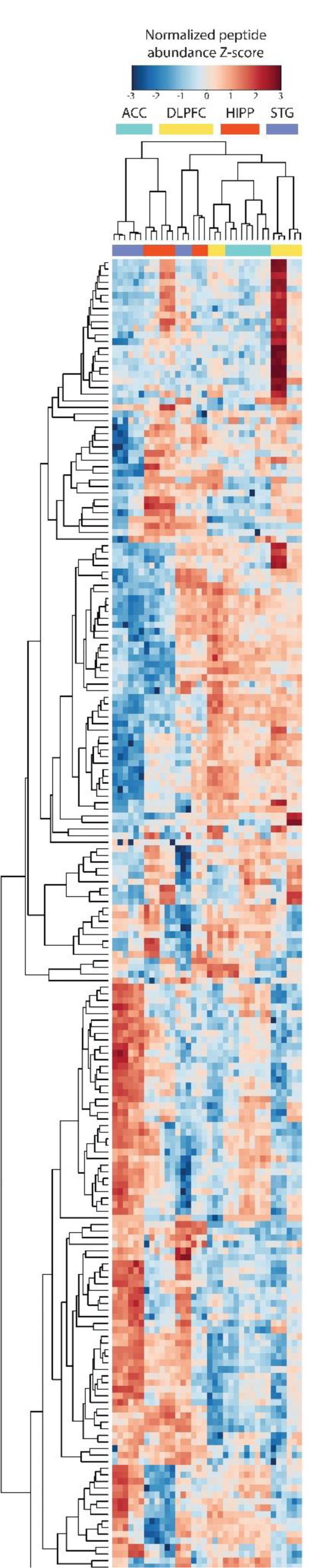
Unsupervised Pearson-correlation cluster analysis. The relative abundance of each peptide was normalized using PSD95 intensity. The data were then Log2-transformed and quantile-normalized. The standard deviation of each peptide was used to rank and select the 200 most variable peptides. Each peptide row was then z-transformed. An unsupervised cluster was created using the *Heatplot* function from the *made4*^*1*^R-package using Pearson correlation.

Because the HPC and ACC gave the most robust clustering results, we focused on these two regions for our subsequent analyses. To derive peptide expression signatures for these two regions, we perfomed a two sample t-test using the geometric mean for each peptide from the three HPC biological samples versus all other regions, followed by the identical analysis of the ACC. We used the resulting value of the t-statistic as a score measuring the degree of differential expression (and thus discrimination) between the HPC and ACC. A scatterplot of t-statistic values for all 200 peptides (ranked based on their t-statistics) shows that approxamitely 50 peptides are expressed at a higher level in the HPC PSD-95 interactome versus the ACC, constituting a putative HPC signature (Figure 5A, red triangles, and Table S1). Conversely, about 50 different peptides are expressed at a higher level in the ACC PSD-95 interactome versus the HPC (Figure 5A, blue squares, Table S1).

**Figure 4.**
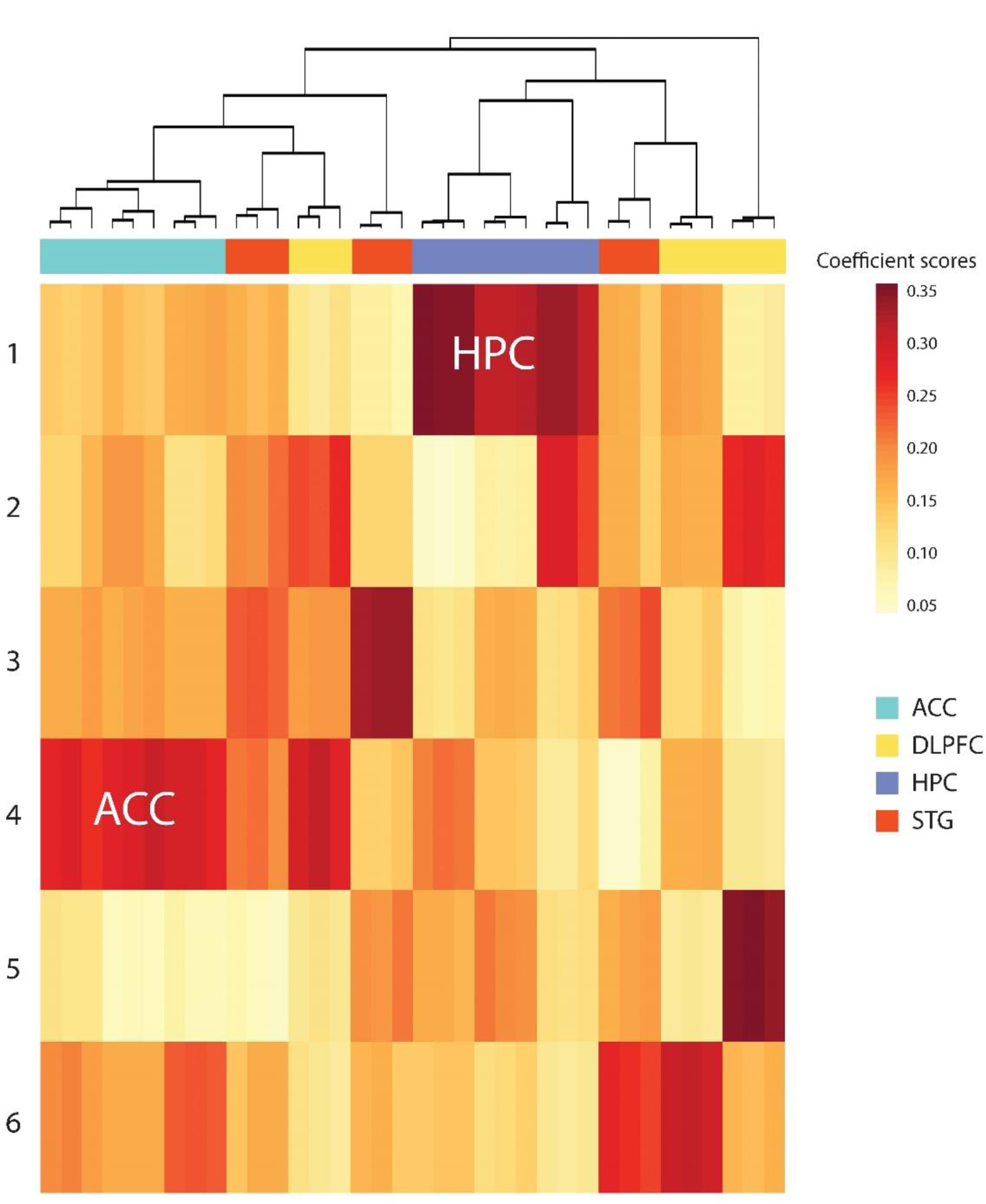
Non-negative matrix factorization (NMF) cluster analysis. Normalized peptide data were analyzed for 200 iterations by least squared minimization. Darker red squares represent high coefficients with 6 vectors, two blocks changed, one for hippocampus (HPC) and another for anterior cingulate cortex (ACC).

**Figure 5.**
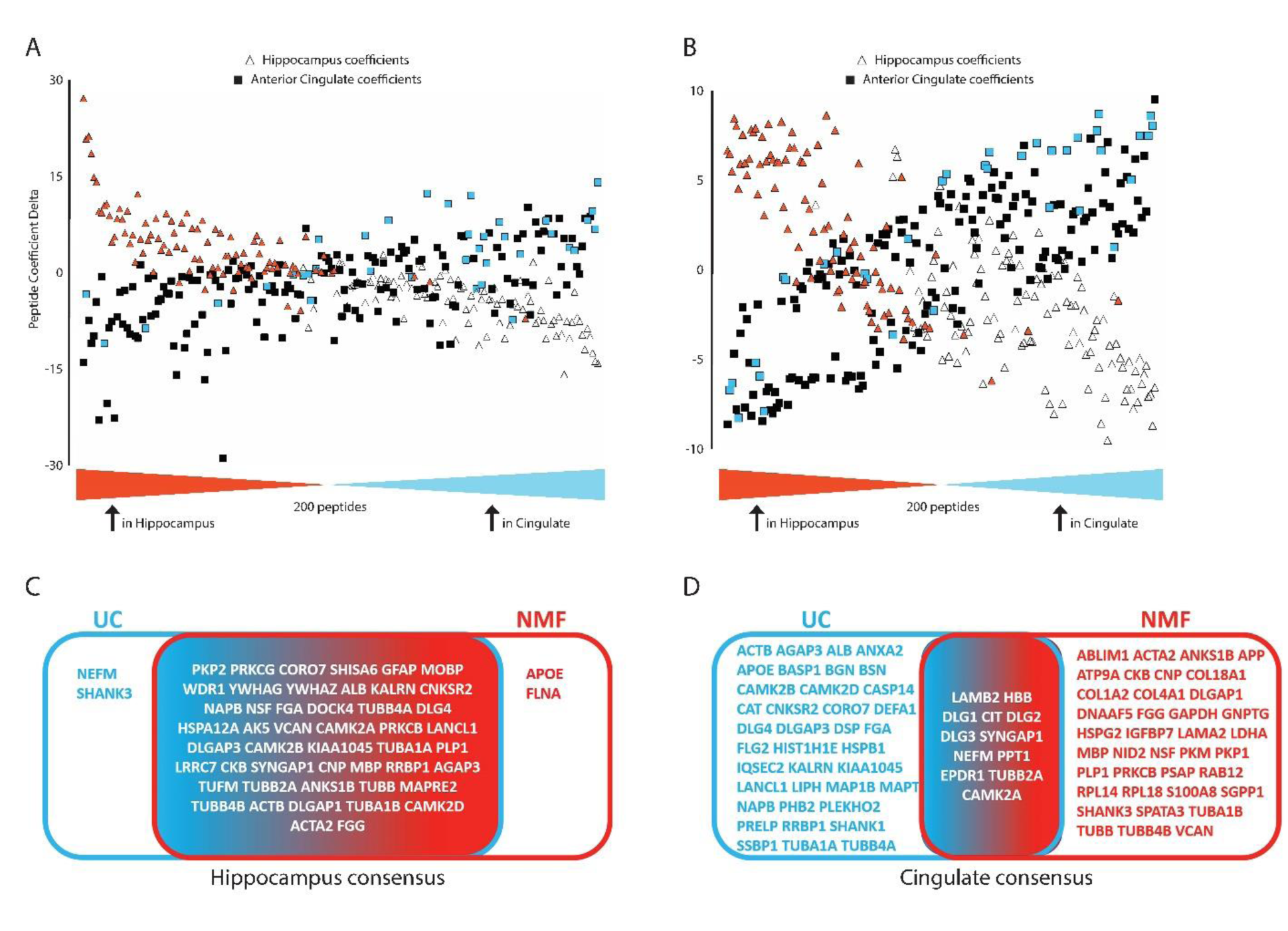
Identification of consensus signatures for the hippocampus (HPC) and anterior cingulate cortex (ACC). Panels A (unsupervised clustering) and B (non-negative matrix (NMF) clustering; semi-supervised) show scatterplots of t-statistic values for all 200 peptides (based on t-statistic rank). 50 peptides are expressed at a higher level in the HPC postsynaptic density (PSD) −95 interactome versus the ACC, identifying a distinct HPC signature for each clustering method (panels A and B, red triangles). Conversely, 50 different peptides were expressed at a higher level in the ACC PSD-95 interactome versus the HPC (panels A and B, blue squares). Peptides found in each signature are listed in table S1. Black squares and open triangles are peptides not included in the peptide signatures from the clustering. Panels C and D show the consensus peptide signatures for the HPC (C) and ACC (D). For the HPC, the 2 clustering methods have significant overlap (46/50 peptides), while the ACC has much less (12/50 peptides).

To validate these findings, we subsequently used an alternative pattern recognition technique to generate peptide signatures which might separate the samples more clearly by reagion. Weakly supervised (by specifying the number of expected vectors) clustering analysis using non-negative matrix factorization (NMF) identified a distinct HPC signature (row 1) when using 6 vectors (Figure 4 and Table S1) [52-54]. Additionaly, the ACC region was also associated with a distinct peptide signature which separated from all other regions (row 4, Figure 4, and Table S1). Next, we employed the same scatterplot comparison approach used above for the signatures from our NMF analysis (Figure 5B). These results also highlight the differential expression of these peptides in the HPC versus the ACC. We found a a high degree of overlap between differential peptide expression and NMF-generated signatures in the HPC (48/50 proteins, Figure 5C), but much lower concordance in the ACC (12/50 proteins, Figure 5D).

Peptide signatures derived from unsupervised clustering, NMF, and differential expression analyses were subsequently integrated into consensus signatures. Specifically, peptide ranks in each signature were combined via geometric mean to generate a consensus peptide signature for the HPC (Table S2) and ACC (Table S3) regions. Multiple peptides belonging to the same protein were combined and the resulting top 25 proteins in the consensus signature for each region were used for bioinformatic analyses to identify pathways, functional categories enriched in each region, and gene knockdown or chemical perturbagen signatures concordant or discordant with our PSD-95 interactome signatures.

We first performed traditional pathway analyses using Toppcluster (Figure 6) [39]. We found overlap of some biological pathways, functions, and molecular disease phenotypes, while others were unique to each region (Figure 6). We also used PiNet (http://pinet-server.org) to explore Panther (Figure 7A) and Kegg (Figure 7B) pathways associated with the HPC signature. Many expected pathways were identified, including glutamate receptor, synaptic transmission, and signal transduction pathways associated with excitatory neurotransmission. Unexpected pathways include inflammatory, immune, and phospholipid-associated functions and processes.

**Figure 6:**
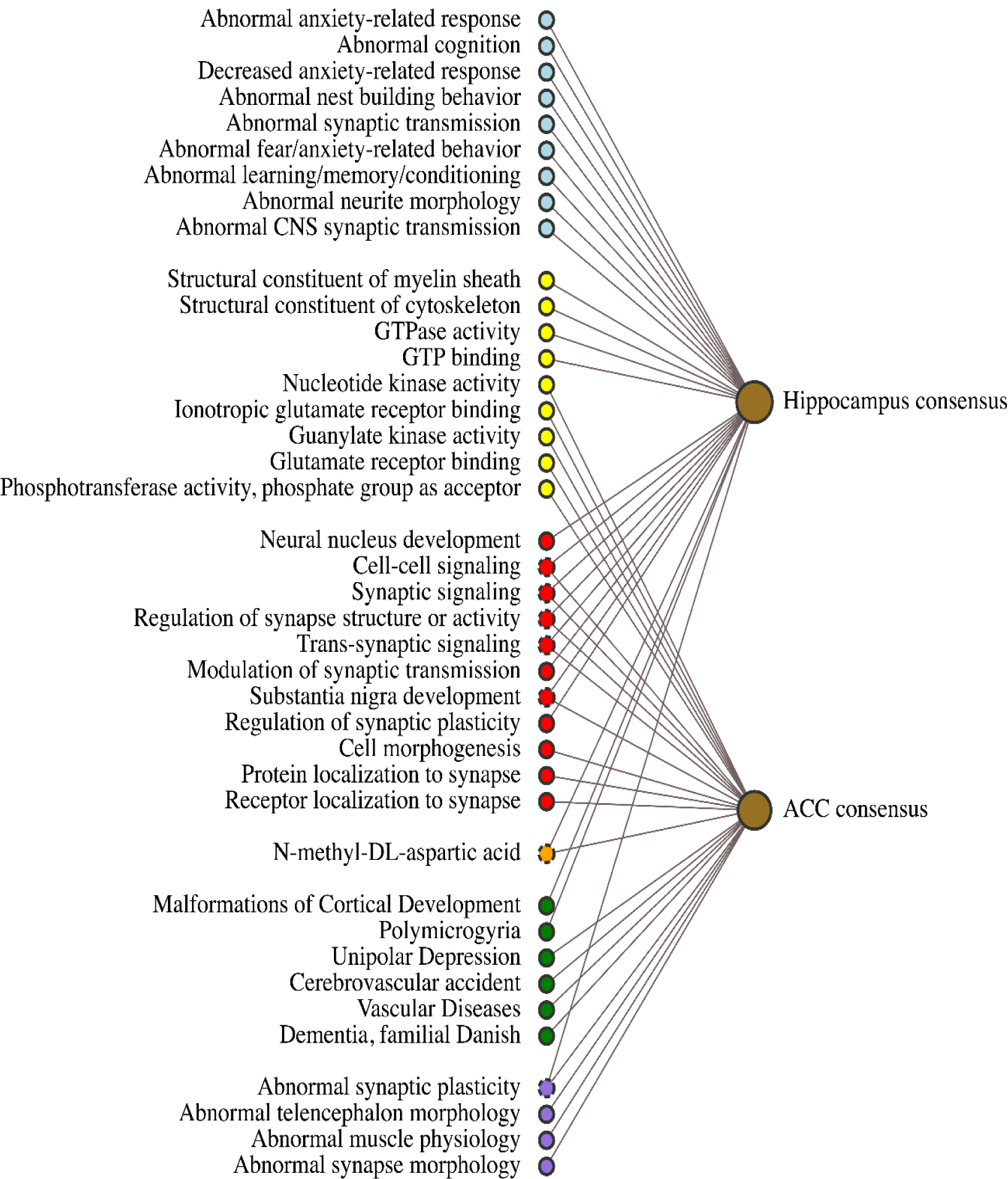
Traditional bioinformatics analyses. ToppCluster Bioinformatic Analysis of Hippocampus and Anterior Cingulate Consensus Signatures. The consesnsus signatures generated after unsupervised and NMF clustering were analyzed by the ToppCluster bioinformatics suite of software. Gene names for the proteins comprising each signature were added as separate clusters, then searched for pathway enrichment. The results displayed represent unique and shared biological functions, pathways, drugs, and phenotypes as determined through curated database mining. The nodes are color coded according to the database/pathway queried: Go: Biological Process (red), Go: Molecular Function (blue), Human Phenotype (purple), Mouse Phenotype (yellow), Disease (green), Drug association (orange).

**Figure 7A.**
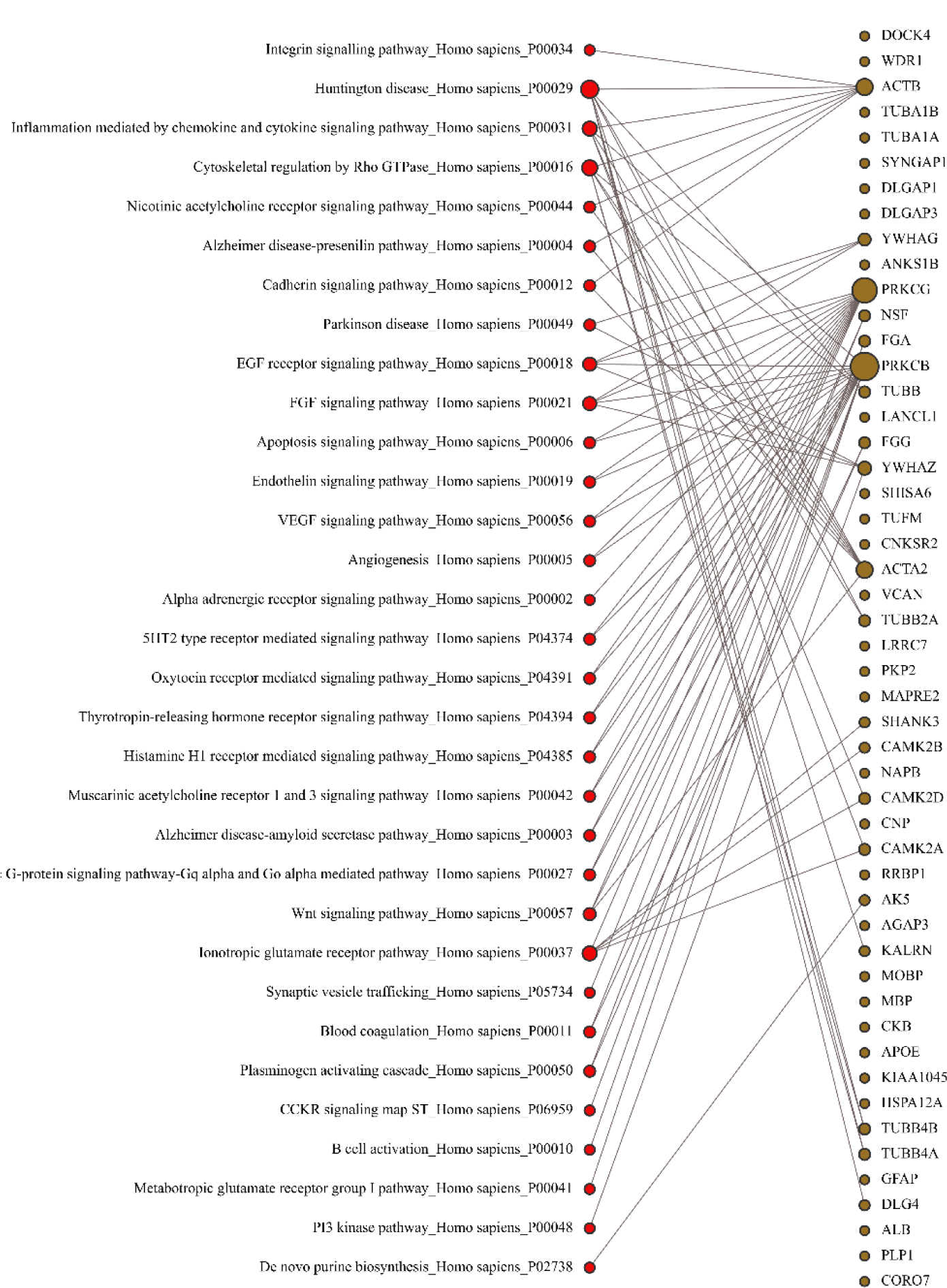
KEGG pathway analysis of hippocampal PSD-95 protein interactome signature. Brown nodes in the figure represent proteins that are over-abundant in hippocampus PSD95 complexes in hippocampus (and thus define the hippocampus signatures). Red nodes represent pathways generated based on clusters of proteins in the hippocampal signature. Specifically, each edge in the figure indicates that a gene encoding a hippocampus signature protein was among differentially expressed proteins for a specific disease or biological pathway, as identified using Enrichr and piNET.

**Figure 7B:**
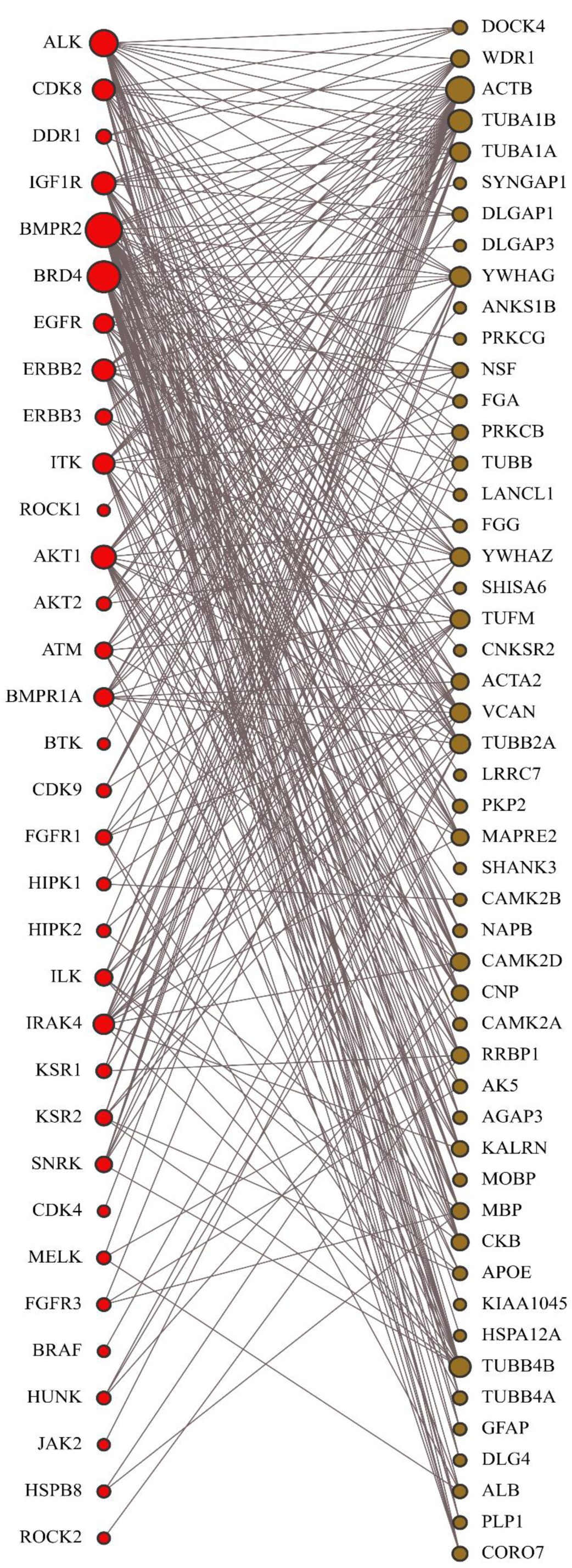
Kinase perturbation analysis of hippocampus PSD-95 protein interactome signature. Brown nodes in the figure represent proteins that are over-abundant in hippocampus PSD95 complexes (and thus define the hippocampus signatures). Red nodes represent kinases whose genetic or chemical perturbation resulted in differential expression of gene sets (effectively defining signatures of these kinases) that overlap with the hippocampus signature. Specifically, each edge in the figure indicates that a gene encoding a hippocampus signature protein was among differentially expressed genes (signature) for a specific kinase, as identified using Enrichr and piNET.

We also used PiNet to explore protein kinase networks associated with our HPC signature (Figure 7C). The BRD4 and BMPR2 kinase pathways have the largest overlap with the HPC signature, followed by ALK, IGF1R, EGFR, ERBB2, CDK8 and AKT1, as indicated by the node radius (or the number of edges). Note also that three of the top five HPC signature proteins (PKP2, PRKCG, MOBP, CORO7, SHISA6), are connected with both BRD4 and BMPR2 pathways. Interestingly, BRD4 is involved in transcriptional regulation of learning and memory, and is expressed in neurons throughout the brain [55, 56]. BMPR2 is expressed in HPC and is implicated in the regulation of anxiety-like behaviors in rodents [57-60].

**Figure 7C.**
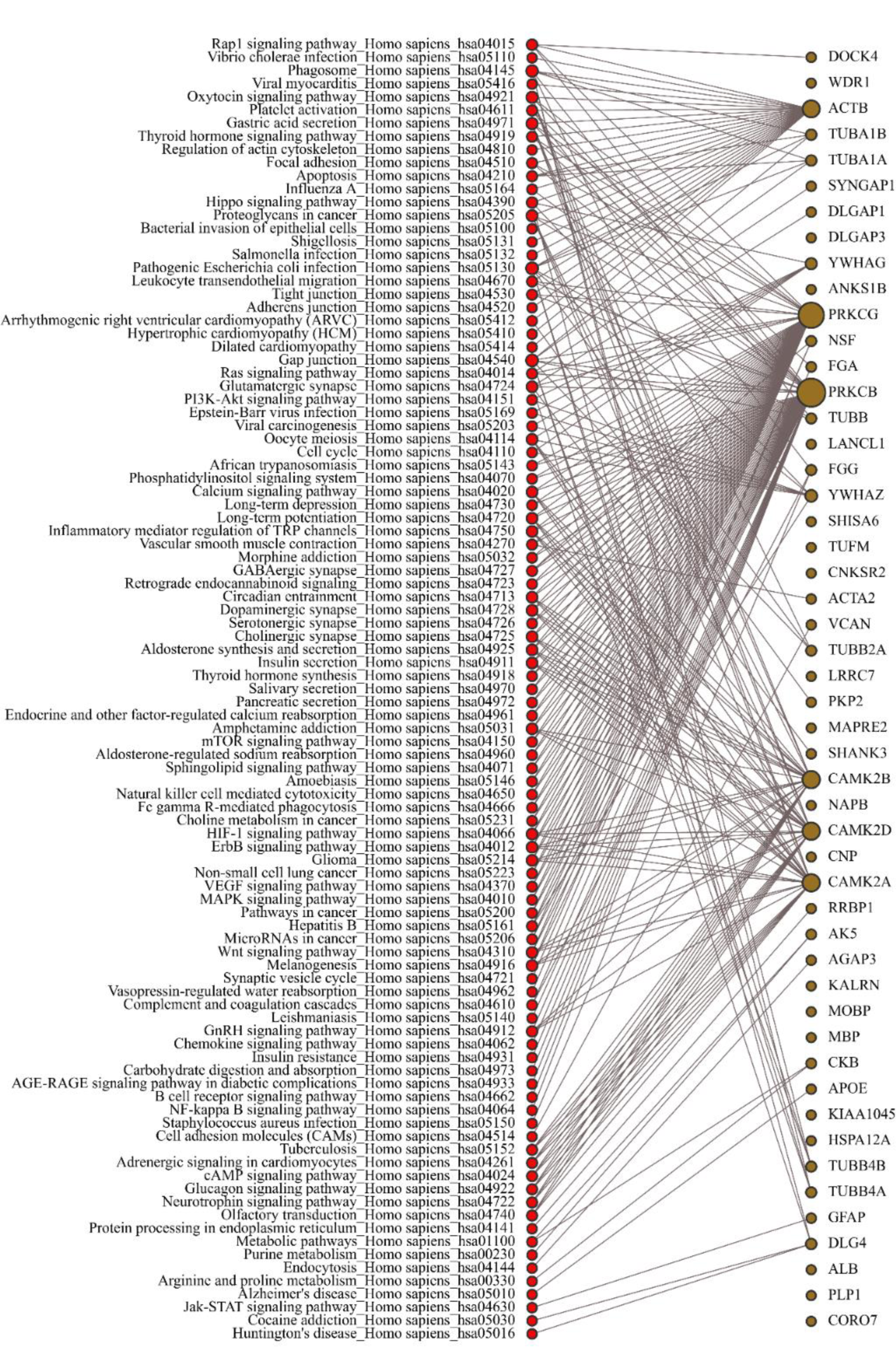
Panther pathway analysis of hippocampal (HPC) PSD-95 protein interactome signature. Brown nodes in the figure represent proteins that are over-abundant in hippocampus PSD95 complexes (and thus define the hippocampus signatures). Red nodes represent biological pathways generated based on clusters of proteins in the HPC signature. Specifically, each edge in the figure indicates that a gene encoding a hippocampal signature protein was among differentially expressed proteins for a specific biological pathway, as identified using Enrichr and piNET.

We next performed exploratory bioinformatics analyses on proteins from our HPC signature using the Library of Network-Based Cellular Signatures (LINCS) database to determine if our signature could help inform functional significance and mechanistic processes. LINCS consists of transcriptional, proteomic, and kinase activity signatures based on chemical (small molecule), genetic (gene knockdown), and kinase activity assays in cellular systems, systematically collected and made available to the community as part of big biomedical data resources [41-43, 45, 46, 61]. Direct search for concordant and discordant LINCS signatures requires a significant probe / protein level overlap. Because of a limited overlap between the HPC consensus signature and L1000 gene set used by LINCS for transcriptional signature collection, we also used an indirect approach consisting of analysis of available gene knockdown signatures contributing significantly to the HPC signature, and identification of small molecules generating discordant (i.e., gene knockdown reversing) signatures (Figure 8A). Such identified small molecules are hypothesized (based on their cell line signatures) to increase expression of genes (and by extension proteins) downregulated in response to a gene knockdown.

**Figure 8A.**
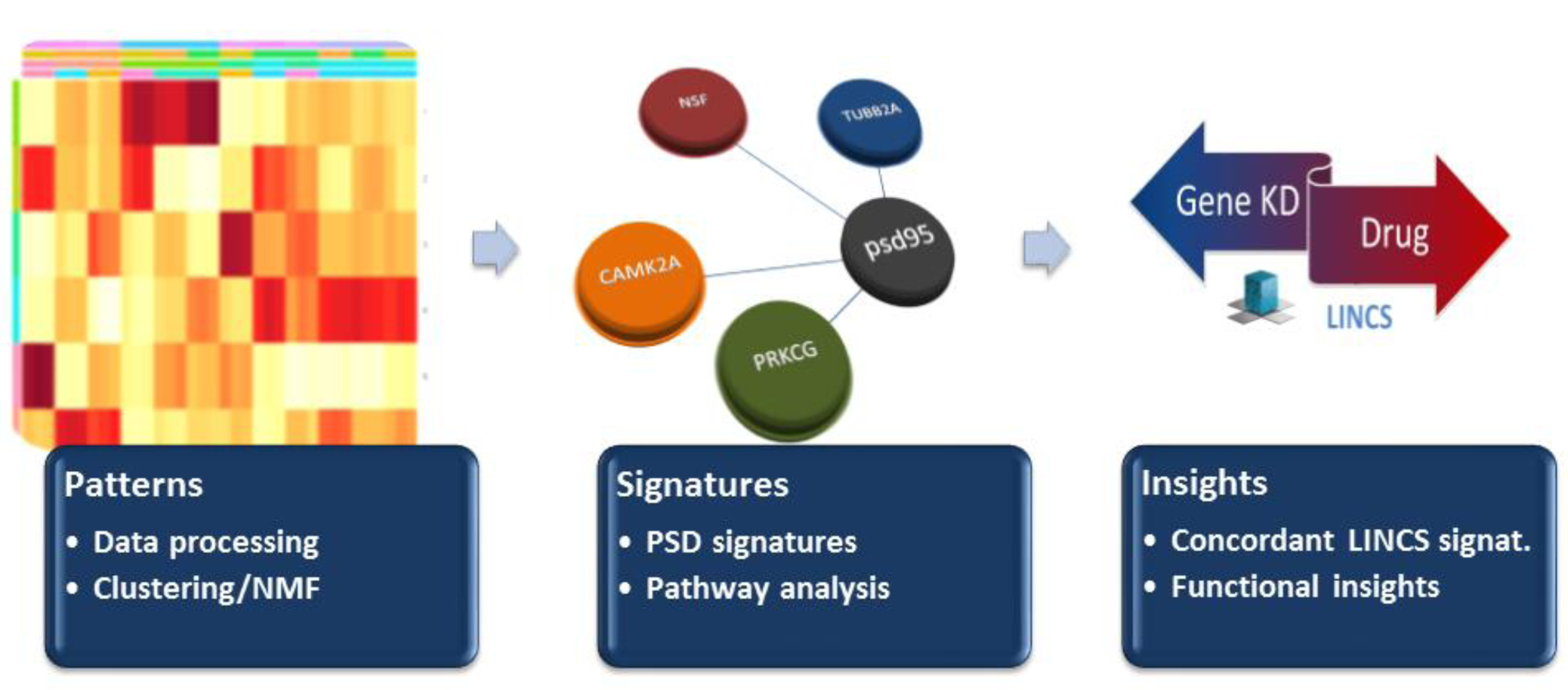
Overview of Library of Network-Based Cellular Signatures (iLINCS) workflow. Following cluster analyses, we used an indirect approach consisting of analysis of available gene knockdown signatures contributing significantly to the hippocampal (HPC) signature. We generated discordant (i.e., gene knockdown reversing) iLINCs L1000 signatures for 18/50 proteins in the HPC signature.

The LINCs database for transcript profiling utilizes mRNAs for the same 972 landmark genes (called L1000). 18 out of 50 proteins in our HPC consensus signature are found in the L-1000 LINCS profile (Table S4). We then searched iLINCS for chemical perturbagen signatures for our 18 genes represented in the LINCs database. This search yielded a list of chemical perturbagens that significantly (adjusted p < 0.05) altered transcript expression for these 18 genes. Interestingly, phenothiazine class drugs increased mRNA expression in the LINCS database for 11 of these targets (Figure 8B). Three of the top five discordant perturbagens belong to this class of antipsychotic medication, including trifluoperazine, thioridazine, and fluphenazine (Figure 8B and Table S4). Other chemicals highly discordant for CaMKIIα iLINCS L-1000 knockdown signature in the VCAP cell line were valproic acid and AG14361. Of note, AG14361 is a potent and selective polyADP-ribose polymerase (PARP-1) inhibitor which has a highly similar chemical structure to Rucaparib, a PARP-1 / ERK2 protein-protein interaction modulator.

**Figure 8B:**
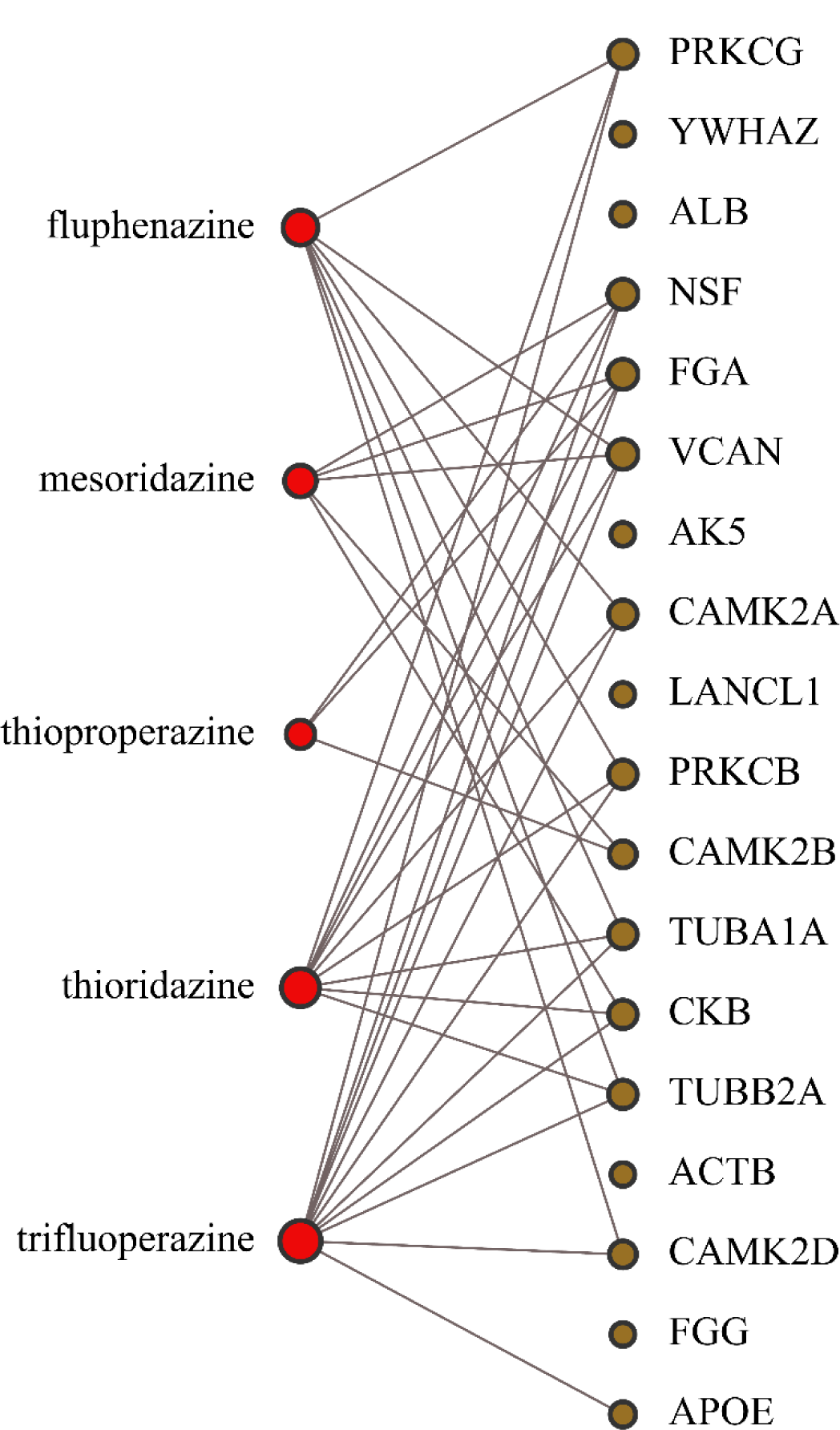
Analysis of correlations between LINCS gene knockdowns for the hippocampus (HPC) PSD-95 protein interactome signature (ie genes that are significantly preferentially associated wih postsynaptic density in HPC) and phenothiazine antipsychotics. Drugs used in analysis (due to their availability in LINCS profiles) are shown as red nodes and include fluphenazine, mesoridazine, thioproperazine, thioridazine, trifluoperazine. The top 50 in genes in the HPC PSD-95 protein interactome signature are represented by brown nodes.

Based on these intriguing findings, we performed an exploratory iLINCS connectivity analysis for HPC consensus signature genes (Table S5). The primary finding from our connectivity analysis was the negative (for lower concentration) correlation of several structurally similar antipsychotics, including thioridazine, trifluoperazine and fluphenazine, with the knock-down signatures of proteins increased in the HPC PSD-95 interactome (Figure 8 and Table S4). These antipsychotics were shown to have low picomolar inhibition of TARP2/ARC protein-protein interactions, suggesting that their binding to Arc protein could represent their primary mode of action [62]. At higher concentration, an inversion to positive correlations is often observed, suggesting that an alternative mechanism could play a role, possibly one that is consistent with antagonism of D2 dopamine receptors by these antipsychotics (albeit with a lower affinity compared to Arc).

## Discussion

Our findings demonstrate that 1) PSD-95 protein hub constituents differ by brain region, 2) we can separate/cluster brain regions based on these differences, and 3) differences in PSD-95 protein hub composion map to divergent functional and pharmalogical substrates. In particular, our results show that the HPC and ACC have unique PSD-95 associated proteomic signatures, in contrast to the DLPFC and STG. This is not surprising given the unique neuroarchitectural features of the HPC with its trisynaptic excitatory circuits, as well as the agranular (ie no layer 4) nature of the ACC [63]. We subsequently focused on the ACC and HPC, generating compartment-specific, proteomic signatures through a pipeline of unsupervised (Figure 3) and semi-supervised NMF clustering (Figure 4), followed by bioinformatic analyses to identify biological pathways. As expected, the most distinct region identified was the hippocampus, which was clearly differentiated via clustering from all other regions tested.

The postsynaptic density and PSD-95 are constitutively modulated through activity dependant mechanisms [14, 18, 64]. Some of the earliest work on characterization of excitatory postsynaptic densities suggested that the composition across regions is relatively uniform [65]. However, our data suggest that there are marked differences in the relative abundance of signaling and other proteins between regions, informing a picture of a non-static environment receptive to immediate and robust action after stimulus. This notion is supported by studies where Peng and colleagues examined the differences between rodent forbrain versus cerebellar excitatory postsynaptic densities, reporting ∼15% of postsynaptic density proteins as significanlty different between regions [10]. Importantly, electrophysiological function of cerebellar synapses are particularly different than cortical synapses. Therefore, we hypothesize that regional functional specificity relies upon inherent differences of postsynaptic protein-protein interactions and that these differences confer functional relevance.

Several distinct approaches have been employed to assess the consituents of the PSD, including fractionation (yielding an enriched fraction of PSDs) [65-71], TAP tagging of PSD-95 for immunoisolation [72], or direct immunoisolation with anti-PSD-95 antibody [48, 73, 74]. Comparison of these different approaches to our present work shows moderate overlap between our human study and the murine flagged PSD-95 dataset (33%), with lower overlap with postsynaptic density fraction studies (between 9-10%) (Tables 2 and S6). The study combining fractionation and PSD-95 isolation was intermediate to these other findings (about 22% overlap with our data). Only one of these studies was performed in humans, and it employed fractionation, making direct comparisons between the datasets of limited value [8]. Other reasons for the varying overlap may include differences in LCMS protocols, normalization procedures, and brain regions analysed. Importantly, there are also differences in control studies. We included an IgG control immunoisolation and qualitatively subtracted proteins from our analyses that were captured with pre-immune IgG. Another source of variation may be our handling of “missing” proteins. We only analysed proteins found in all four brains regions. Because missing proteins may either be true negatives, and represent important differences between regions, or be proteins that did not have detectable fragments in the LCMS protocol (false negatives), we may have eliminated several region-specific proteins that may be found in the PSD and/or PSD-95 interactome.

**Table 2.**
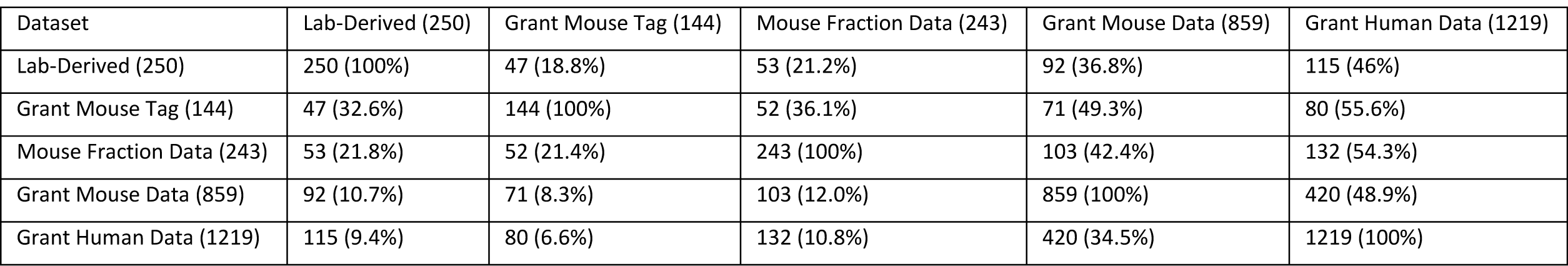
Overlap of PSD-95 interactome proteins vs selected datasets. Percentages in reference to rows.

Regarding the brain regions employed, the mouse studies included in Table 2 and S6 all used whole brains as input, while we used discrete brain regions [72-76]. The human postsynaptic density fractionation study combined frontal, temporal, parietal, and occipital brain regions [77]. The differences in species and brain region notwithstanding, these data suggest that the PSD-95 interactome represents about 20% of the postsynaptic density-compartment’s proteome. This is not surprising given the volume and complexity of this subcellular compartment. Further work is needed to make more definite conclusions regarding similarities or differences across species and to account for divergences associated with specific methodologies.

In contrast to the aforementioned murine studies, we assessed individual brain regions, an approach made feasible by the relative abundance of human brain compared to rodents. We compared the results of unsupervised and NMF clustering signatures from HPC and ACC. The HPC consensus signature (Figure 5C and Table S2) shows nearly complete overlap (48/50 proteins) between the two clustering methods, suggesting that the HPC signature is clearly identifiable and unique from the other regions tested. In contrast, the ACC signatures had less overlap, with 12/50 proteins common to the unsupervised and NMF signatures (Figure 5D and Table S3). This is likely due to structural and functional similarities between the ACC, DLPFC, and STG, which are all regions of input processing and integration for higher order cognitive function [77]. Between the two clustering methods, the NMF analyses appears to better differentiate the HPC and ACC compared to the unsupervised hierarchical clustering approach. NMF is a semi-supervised analysis which relies on the user to select the number of vectors or groups the data should fall into. Each vector represents a linear metagene comprising all the data in the set. The dimentionality of metagenes are reduced through decomposition into two nonnegative matrices, then matched back to the metagene through an iterative multiplicative updates algorithm (in this study we used 200 iterations for all NMF analyses) [28].

Nonnegative matrix factorization (NMF) was introduced as a weakly supervised approach to analysis and interpretation of large data sets, including molecular pattern discovery, class comparison and prediction in computational biology. In particular, NMF has been increasingly popular as a method of choice for omics data consisting of abundance levels for tens of thousands of genes or proteins across a number of samples. In this context, NMF involves the decomposition of a nonnegative protein (or peptide) expression matrix *P* consisting of observations on *p* peptides from *n* samples, into two nonnegative matrices *W* and *H*. Each column of *W* defines a meta-protein (defined as a linear combination of individual proteins or peptides that can be viewed as a signature of some underlying biological state with proteins obtaining larger coefficients contributing more to that state), while each column of *H* represents the meta-protein expression pattern of the corresponding sample. NMF represents an alternative to other pattern recognition and data mining techniques, such as unsupervised hierarchical clustering or Principal Component Analysis, yielding results that are largely comparable but often easier to interpret, in particular as it pertains to the derivation of molecular signatures of biological states. Here, these signatures are defined by peptides whose high expression is associated with specific brain regions, as reflected by those peptides obtaining high coefficients in the corresponding meta-protein columns of *W* [28]. In summary, we conclude that combining the unsupervised and NMF approaches appears to identify concensus clusters, providing robust results for our our PSD-95 affinity purification-LCMS protocol.

Using the consensus signatures generated via combining consensus hits from NMF and hierarchical clustering, we performed pathway analyses using ToppCluster (Figure 6) [39]. Top pathways for the HPC include GTP binding, GTPase activity, and cytoskeletal regulation, while top pathways for the ACC include glutamate receptor binding and nucleotide kinase activity (Figure 6). Interestingly, the top hits from the HPC signature reflect fundamental biological processes that are prominent in this region, including a high density of exciatory synapses with excitatory (ie glutamate) inputs. For example, SHISA6, PKP2, PRKCG, NSF and CORO7 are important components of postsynaptic activity, memory, and learning. Consistent with the biological functions of these proteins, behavioral phenotypes of the HPC include abnormal learning and memory, anxiety, and nest building using targeted knodckdown or deletion of these genes [78-84].

Given the structural and developmental differences between the HPC and ACC, our findings suggest that the unique consensus signatures represent individualized structural and behavioral consequenses in a region-dependent manner. Consideration of specific elements of each signature may inform the overall fidelity of our approach. For example, SHISA6 is an auxillary AMPA-subtype glutamate receptor binding protein important for trafficking in association with PSD-95 [85]. SHISA6 contains a PDZ domain and directly binds to PSD-95 [85]. Expression of SHISA6 in HEK cells altered AMPA receptor GluA1 and GluA2 synaptic localization and function. Additionally, SHISA6 expression is preferentially expressed in all hippocampal subfields and the dentate gyrus, but has limited expression in the cortex [85, 86]. Identification of this molecular componenet of neuroplastic functions attributable to the HPC suggests that the HPC signature is in fact very “hippocampal.”

Several other proteins in our HPC signature, including PRKKCG, NSF, βSNAP, PICK1, and PKC, also have strong ties to excitatory neuroplasticity. Initial studies on PRKCG indicated deficits in spatial and contectual learning, likely due to abnormal hippocampal LTP induction in mutant PRKCG mice [87, 88]. Others have shown that PKC gamma traffics AMPARs in zebrafish in an NSF and PICK1 dependent manner, resulting in miniEPSC potentiation [89-91]. Additionally, NSF attachment protein beta (NAPB or β-SNAP) can regulate NSF and PICK1 mediated AMPAR trafficking, where βSNAP directly binds to PICK1 and allows NSF to dissasemble PICK1-AMPAR complexes [92]. NSF and βSNAP are highly expressed in hippocampal neurons and regulate AMPAR recycling and internalization in an NMDAR activity and Ca^2+^ dependent manner [93-98].

Other proteins in the HPC signature have previously been demonstrated as highly specific to this region. For example, plakophilin 2 (PKP2) represents our most robust contributor to hippocampal specific signature, although not much is known about how it functions at synapses. PKP2 belongs to the armadillo-repeat and plakophilin family of genes including β-catenin, Sys1, Plakophilin-1, Plakoglobin, and Importin-α among many others [99]. One study showed PKP2 transcript expression was highly specific for hippocampus/Ammon’s horn in mouse brain [100]. The presence of these previously identified HPC-specific proteins supports the conclusion that our signature differentiates the HPC from other brain regions.

Another biological theme associated with the HPC signature relates to structural molecules that facilitate clustering of signaling molecules to the PSD. LRRC7 (also known as Densin-180) is an ERBIN (Erbb2 interacting protein) paralog which localizes to PSDs and directly interacts with CaMKIIα and α-actinin [101-105]. Densin-180 also interacts with maguin-1, more commonly referred to as connector enhancer kinase of ras 2 (CNKSR2) or CNK homolog protein 2, which mediates Densin-180/maguin-1 complex formation with PSD-95 at glutamatergic synapses [106-108]. Changes in Densin-180 expression impact dendritic branching, localization of PSD95 to synapses, and clustering with shank3 and other proteins [107-111]. Alterations in other structural proteins found in the HPC signature, including delta-catenin, are associated with developmental disorders, including autism and cri-du-chat syndrome [112-117]. Post-translational modifications appear to mediate clustering of catenins in the PSD; palmitoylation of beta-catenin by DHHC5 increases N-cadherin/beta-catenin interaction, AMPAR insertion into the plasma membrane, and increased spine volume [118, 119]. In summary, many of the proteins in the HPC signature have higher or exclusive expression in the HPC, as well as serve critical roles in neuroplastic functions that underlie the functional novelty of this region [120].

In contrast to the HPC, the ACC is not as extensively studied, and thus there is a much smaller literature to draw from regarding ACC-specific expression and function of specific proteins found in this signature. In addition, the ACC consensus signature has fewer hits than the HPC, likely due to overlap of its PSD contents with the more closely functionally related DLPFC and STG, which are also neocortical regions. Interestingly, the ACC has an important gross structural dissimilarity compared to DLPFC and STG, in that it lacks a granular layer (it is considered agranular cortex with no formal layer IV) [63].

A family of molecules that facilitate protein-protein interactions and clustering with ionotropic glutamate receptors is overrepresented in the ACC signature [121]. DLG1 (aka SAP97), DLG2 (PSD93 or chapsyn-110), and DLG3 (SAP102) are highly homologous and have similar functions, likely the result of gene duplication. Despite their sequence similarity, they possess unique sequences/motifs that promote diverse protein-protein interactions [122, 123]. Alterations in these glutamate receptor interacting proteins have been found in various brain disorders, including schizophrenia and dementia [124-127], suggesting a unique role for the ACC in the pathophysiology of cognitive dysfunction in severe neuropsychiatric illness. This notion is supported by the cognitive functions ascribed to this limbic region, which include working memory and executive function [128] [129].

Another informative hit in the ACC signature is LAMB2, a protein that regulates postsynaptic synapse formation [130, 131]. Null mutations for LAMB2 yield disorganized cortical lamina, and its localization specifically to limbic cortex suggests involvement in the biology of higher cortical functions [132]. Another hit with biological relevance is CIT (Citron Rho-Interacting kinase), which directly binds to PSD-95 via the PDZ3 domain [133, 134]. CIT expression in HPC is relatively low (and only in GABAergic interneurons), whereas cortical expression is more robust and not limited to interneurons [103, 135]. CIT is a kinase which phosphorylates RhoA and RhoB, small GTPases which regulate actin cytoskeleton dynamics. [136-139].

The identification of proteomic signatures that uniquely represent specific brain regions suggests an important opportunity for elucidation of pathophysiology mechanisms as well as identification of pharmacological agents which may simulate or reverse pathological changes found in disease states. One powerful tool developed to facilitate such analyses involves transcriptional profiling and connectivity analyses with large databases of altered biological substrates. We employed the LINCS database to develop insights into the observed region-specific variation in PSD95 postsynatic complexes. Using recently developed iLINCS software (http://www.ilincs.org), we identified small molecules that generated either discordant (i.e., gene knock-down reversing) or concordant (resulting in similar transcriptional echo to that of gene knock-down) signatures from our studies. Interestingly, 12 out of 18 gene knock-down signatures included in the analysis, including PRKCB, CAMK2A and TUBB2A, are “reversed” by phenothiazines including trifluoperazine, thioridazine and fluphenazine (Figure 8 and Table S4). This class of D2 receptor antagonists promotes a “hippocampal” pattern of gene expression, resotring a more HPC-like profile of transcript expression in cell culture.

This observation was suprising as these drugs are well-characterized as D2 dopamine receptor antagonists, while most of the hits in our signatures were related to excitatory (glutamatergic) neurotransmission [140]. Since dopamine is a potent regulator of glutamate transmission, we would have expected to see more dopamine associated proteins in our signatures [141-144]. However, recent work puts our findings into a different context. A new mode of action was recently reported for these drugs that involves binding and regulating PSD proteins, including Arc and TARPγ2, regulators of synaptic activity in hippocampus [62]. Interestingly, thioridazine and trifluoperazine affinity for Arc was higher than for D2 receptors, raising the possibility that dopaminergic modulation may be secondary to effects on PSD constituents. While further work is needed to explore this putative alternative mechanism, it raises intriguing possibilities for modulating HPC related functions, such as learning and memory.

In addition to this initial observation regarding phenothiazines and our HPC signature, the LINCS database includes varying concentrations of drugs. The gene knock-down signatures discussed above were only restored by phenothiazine drugs at lower concentrations, while concordant responses (ie simulating knockdown or loss of critical HPC-specific gene expression patterns) were observed at higher concentrations and longer exposure times (Table S5). This finding suggests that effects seen at lower doses may be secondary to targets of higher binding efficiency, including the PSD95 interactome constituents ARC and TARPγ2.

Other findings from our iLINCS analyses HPC-like transcriptional signatures may be pharmacologically modulated. For example, we found a strongly discordant (to the HPC) signature for PARP1 inhibitor analog, AG14361. Recent evidence shows inhibition of PARP1 causes loss of LTP induction in rodents [145, 146]. Disruption of PARP1-ERK2 protein-protein interactions in the nucleus were responsible for this loss [145, 146]. Our bioinformatic analyses suggests that synaptic protein-protein interactions with CaMKIIα, CaMKIIβ, PRKCG, or NSF may be disrupted by PARP1 inhibitors. This may be a result of reduced transcription of these targets in a PARP1 dependent manner. In summary, the signature-based, LINCS analyses suggest some interesting mechanistic clues, provided additional support for the recent discovery of a new mechanism or action for phenothiazine antipsychotic drugs, and spotlight specific experimental steps for further validation and potential drug repositioning.

The samples, process, and methods used in this study aren’t without certain limitations. For example, we used only 3 biological replicates, all male, and elderly (71-73 y.o.). Using elderly subjects could inherently skew the data, as synaptic protein-protein interactions most likely change as we age. Further studies using both males and females of a variety of ages would increase the significance and application of these findings. Also, transient or low abundant interactions in the PSD-95 interactome are likely missed due to limit of detection issues with quantitative mass spectrometry. Finally, we only analyzed 4 brain regions, while isolating PSD-95 from more diverse brain regions may allow for greater fidelity of low abundant interactions.

Another caveat for our study is the lack of availability of neuronal cell lines for our bioinformatics studies. Several neuronal cell lines that would be expected to constitute a better proxy for brain tissue, including NEU and NEU.KCL, are included in the current version of the LINCS L1000 signature library. Unfortunately, none of the genes whose products are preferentially associated with PSD-95 post-synaptic protein complexes in hyppocampus have shRNA knockdowns in these neuronal cell lines at this time. However, a number of anti-psychotic drugs identified here as reversing KD signatures of those genes in other cell lines, such as VCAP, have also been profiled in neuronal cell lines. Their signatures are in general positively and significantly correlated with those of other related anti-psychotics, both in NEU and VCAP cell lines. For example, trifluoperazine NEU signature (LINCSCP_41445) yields a correlation (when using top 100 L1000 genes) of 0.41 with trifluoperazine VCAP signature (LINCSCP_64601), 0.42 with thioridazine VCAP signature, and 0.40 with fluphenazine VCAP signature, respectively. The negative correlations with gene knock-downs also extend to NEU anti-psychotic signatures, with trifluoperazine NEU signature resulting in correlations of −0.32 (p-value of 0.001), −0.34 (p-value of 5.0E-04), and −0.23 (p-value of 0.02) for VCAN, PRKCB, and TUBB2A gene knock-downs in VCAP cell lines. Thus, while direct evidence of correlations between anti-psychotic and gene KDs signatures in neuronal cell lines is lacking at present, based on these comparisons it is likely they persist.

## Acknowledgments

We would like to thank all those that have supported this research: the L.I.F.E. Foundation, UCNI pilot award, Lindsay Brinkmeyer Schizophrenia Research Fund, Center for Clinical and Translational Science Training (CCTST), and NIH grant MH107916. The MS data was acquired in the University of Cincinnati Proteomics Laboratory on a mass spectrometer funded in part through the NIH S10 shared instrumentation grant RR027015.

---

**Table S1.**
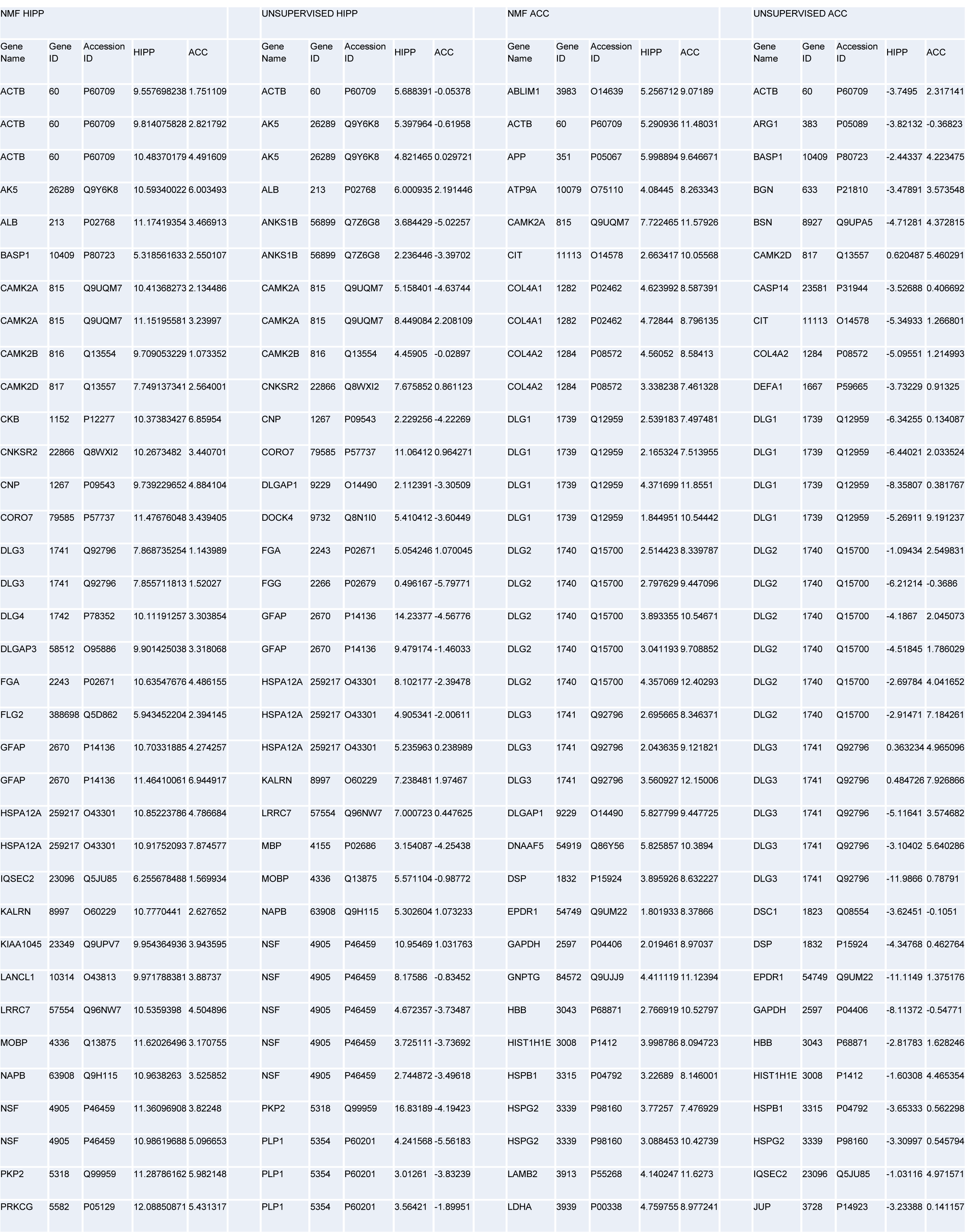
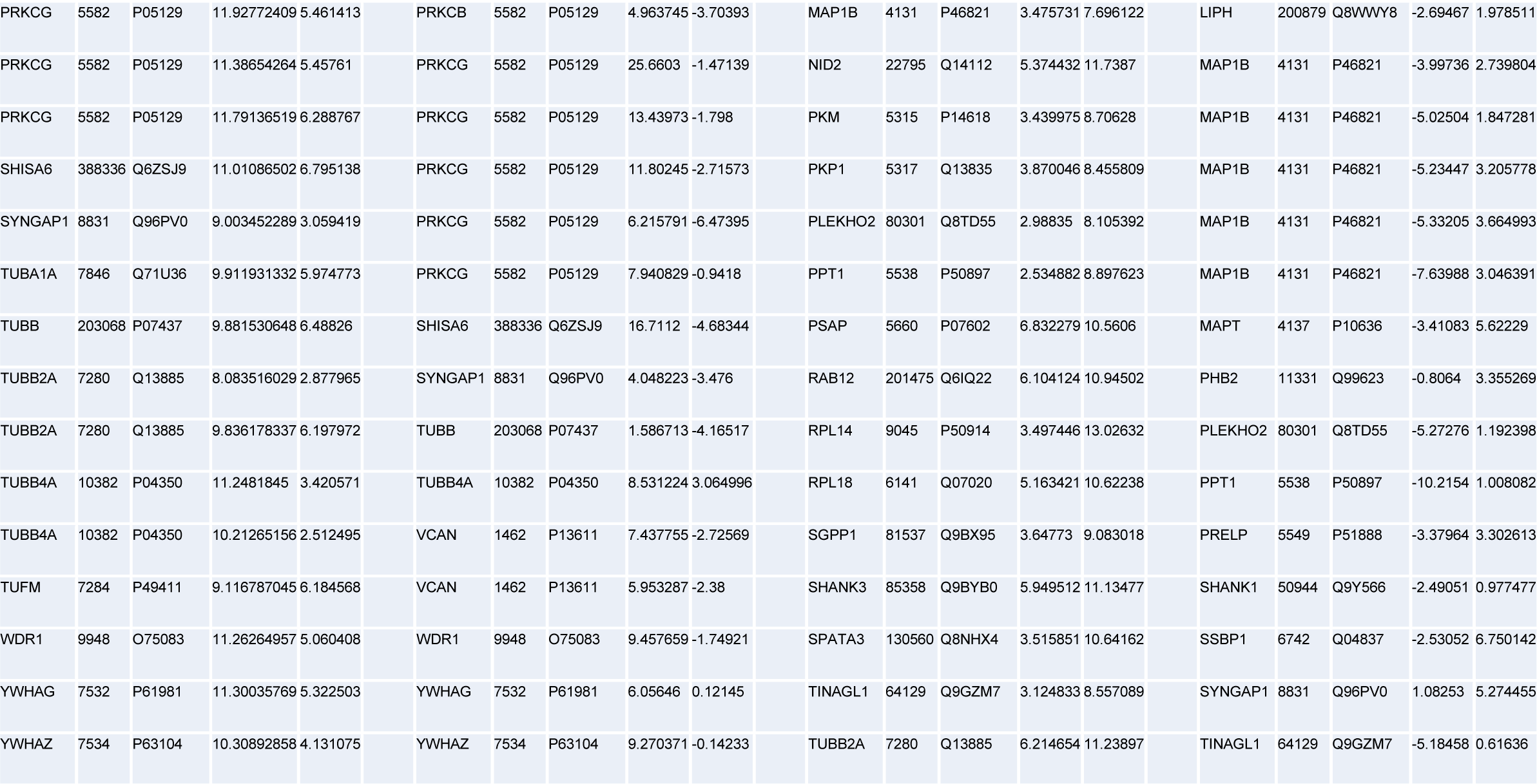
Hippocampus and ACC protein signatures from unsupervised and NMF clustering analyses.

**Table S2.**
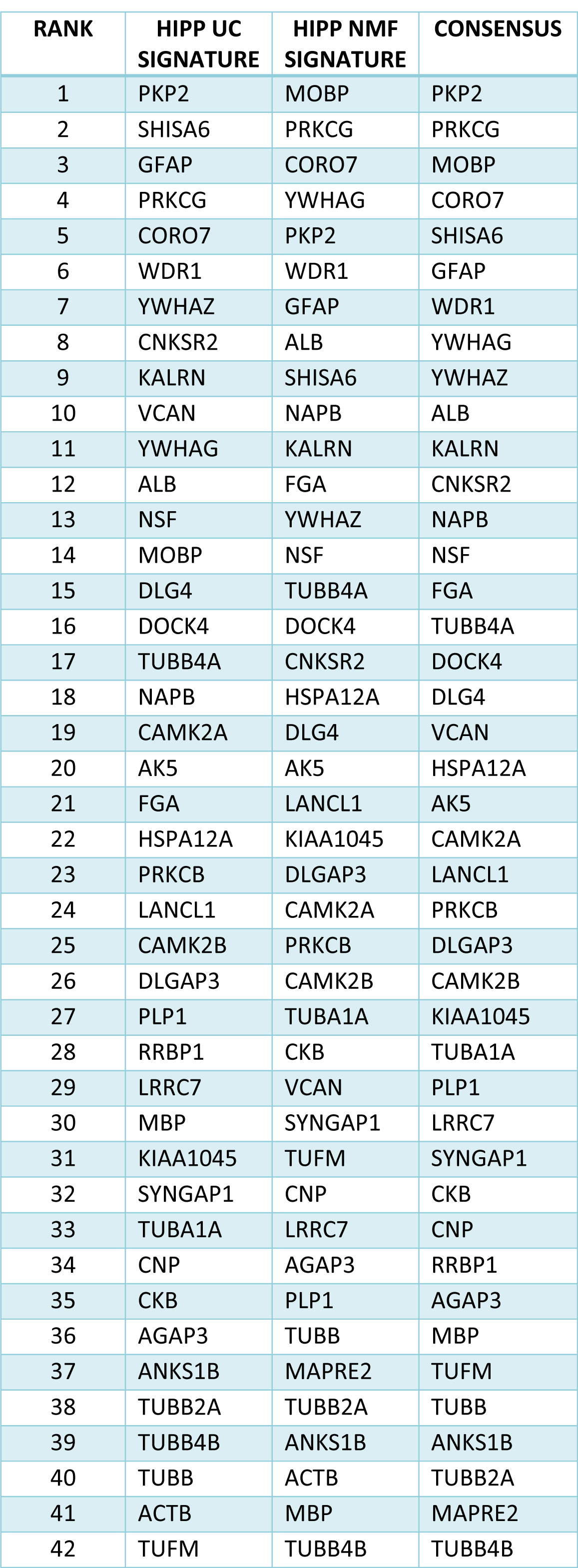
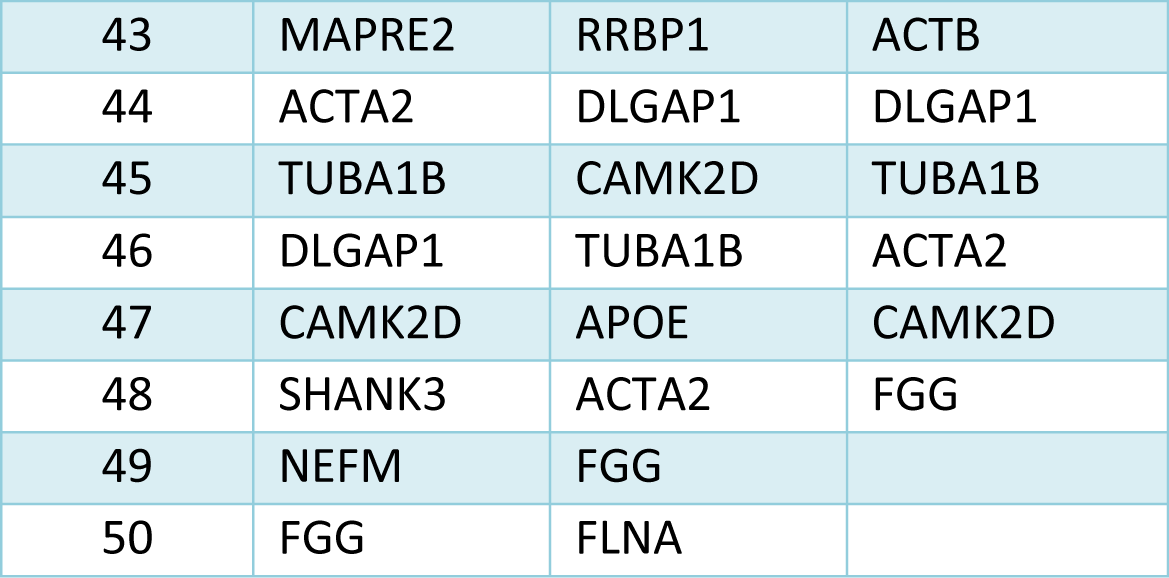
Concesus signature for the hippocampus.

**Table S3.**
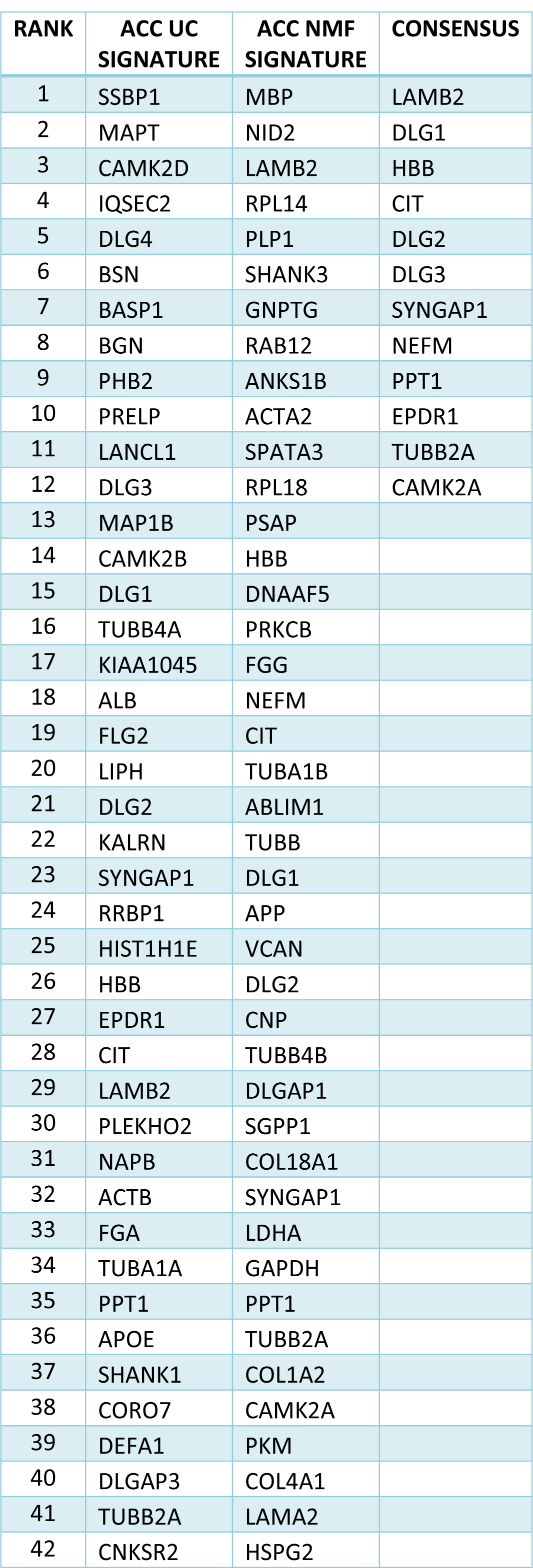
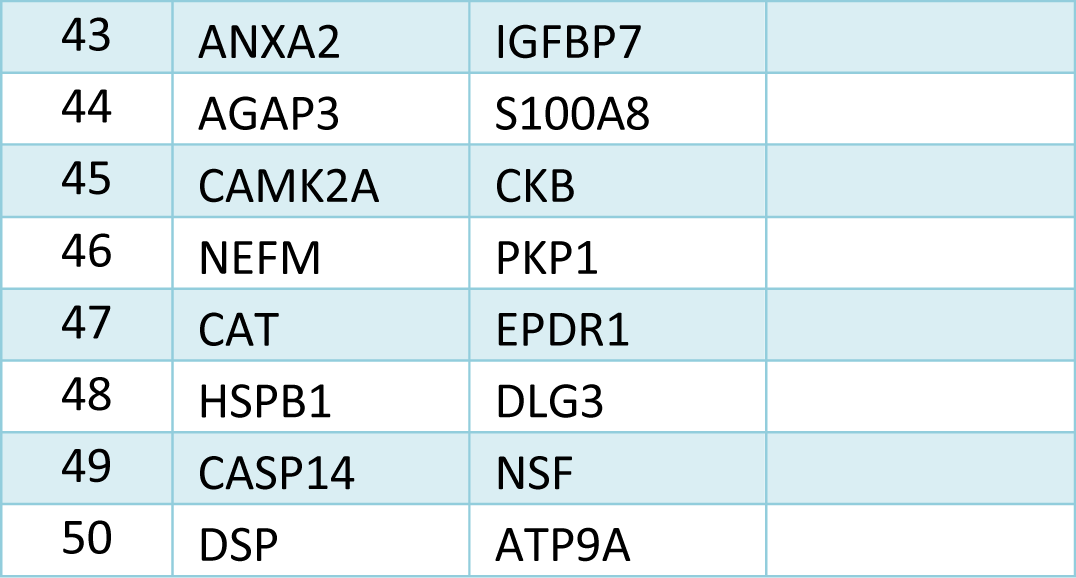
Concesus signature for the anterior cingulate cortex.

**Table S4.**
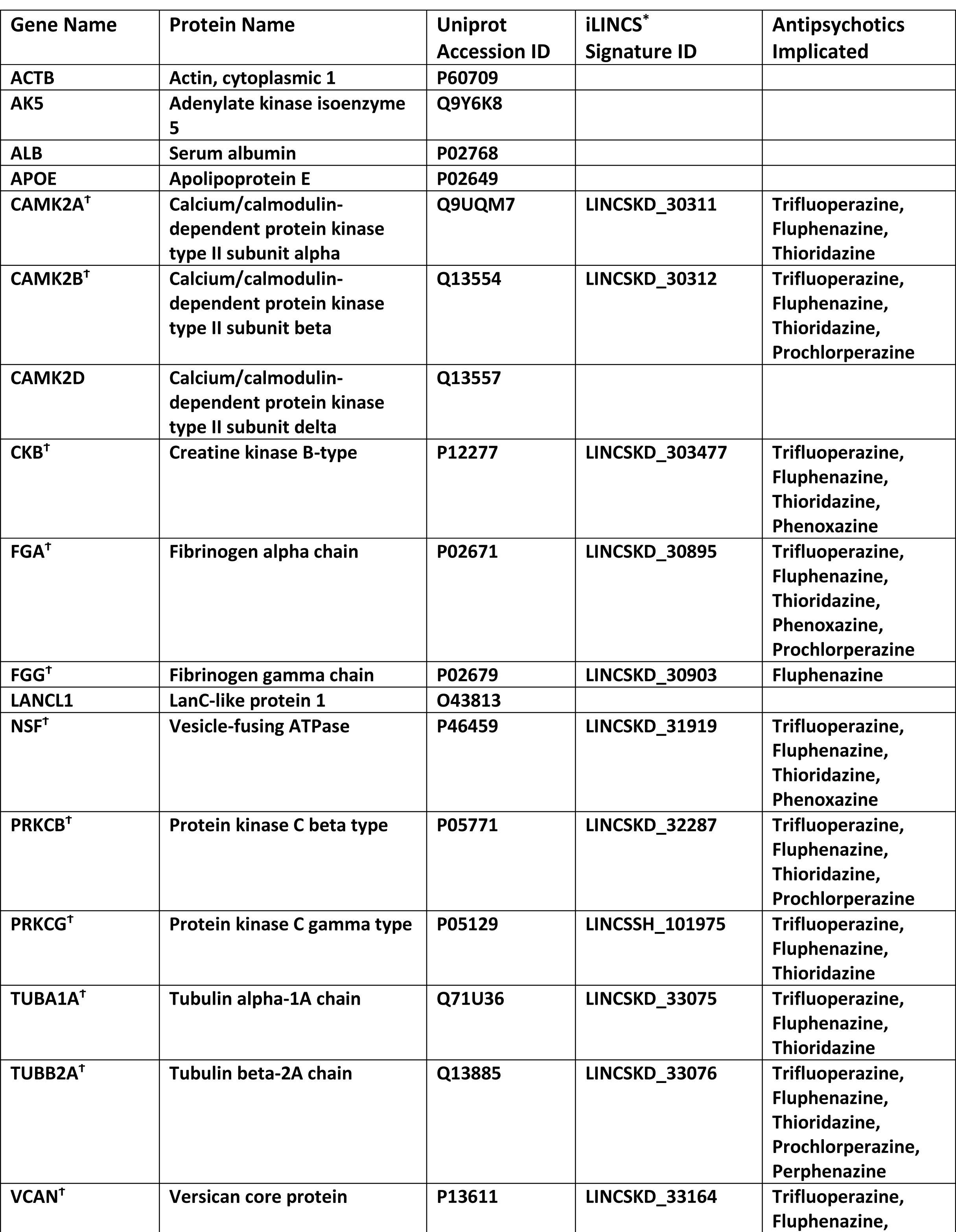
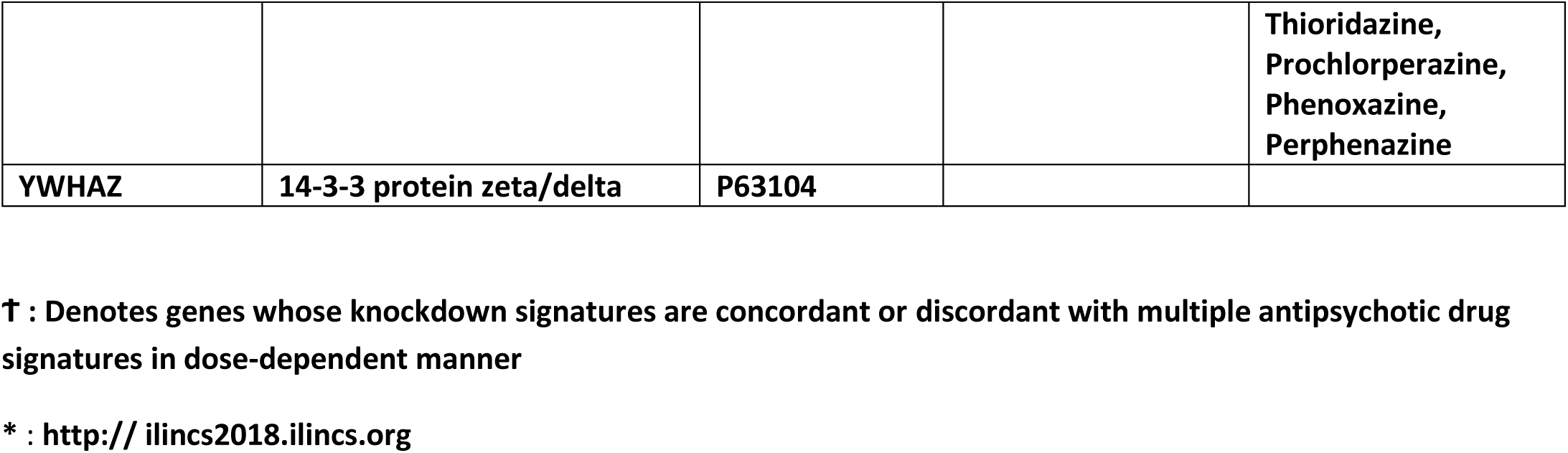
Hippocampus consensus signature iLINCS analysis.

**Table S5.**
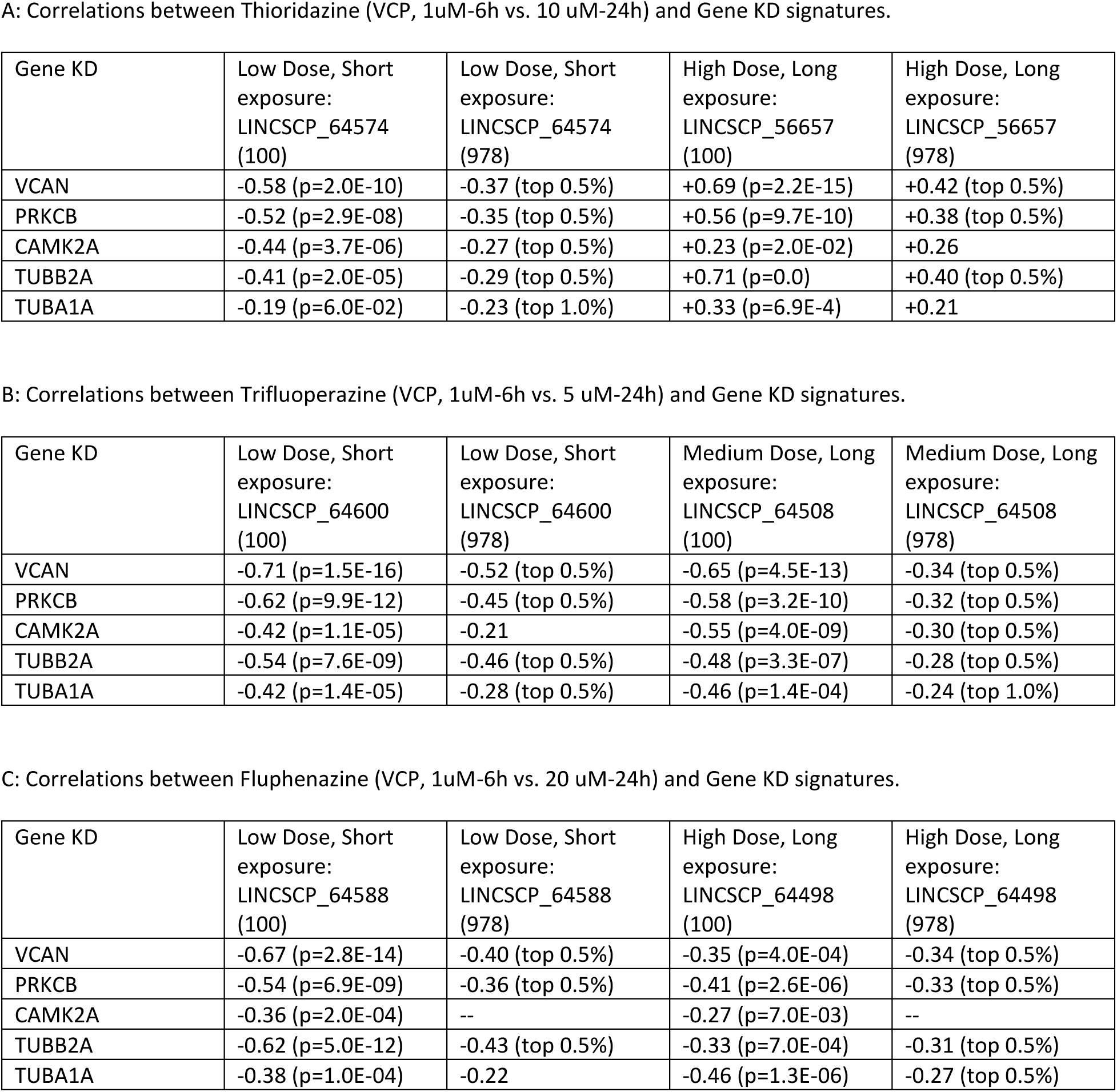
iLINCS connectivity analyses.

A: Correlations between Thioridazine (VCP, 1uM-6h vs. 10 uM-24h) and Gene KD signatures.
| Gene KD | Low Dose, Short exposure:<br>LINCSCP_64574<br>(100) | Low Dose, Short exposure:<br>LINCSCP_64574<br>(978) | High Dose, Long exposure:<br>LINCSCP_56657<br>(100) | High Dose, Long exposure:<br>LINCSCP_56657<br>(978) |
| --- | --- | --- | --- | --- |
| VCAN | -0.58 (p=2.0E-10) | -0.37 (top 0.5%) | +0.69 (p=2.2E-15) | +0.42 (top 0.5%) |
| PRKCB | -0.52 (p=2.9E-08) | -0.35 (top 0.5%) | +0.56 (p=9.7E-10) | +0.38 (top 0.5%) |
| CAMK2A | -0.44 (p=3.7E-06) | -0.27 (top 0.5%) | +0.23 (p=2.0E-02) | +0.26 |
| TUBB2A | -0.41 (p=2.0E-05) | -0.29 (top 0.5%) | +0.71 (p=0.0) | +0.40 (top 0.5%) |
| TUBA1A | -0.19 (p=6.0E-02) | -0.23 (top 1.0%) | +0.33 (p=6.9E-4) | +0.21 |

B: Correlations between Trifluoperazine (VCP, 1uM-6h vs. 5 uM-24h) and Gene KD signatures.
| Gene KD | Low Dose, Short exposure:<br>LINCSCP_64600<br>(100) | Low Dose, Short exposure:<br>LINCSCP_64600<br>(978) | Medium Dose, Long exposure:<br>LINCSCP_64508<br>(100) | Medium Dose, Long exposure:<br>LINCSCP_64508<br>(978) |
| --- | --- | --- | --- | --- |
| VCAN | -0.71 (p=1.5E-16) | -0.52 (top 0.5%) | -0.65 (p=4.5E-13) | -0.34 (top 0.5%) |
| PRKCB | -0.62 (p=9.9E-12) | -0.45 (top 0.5%) | -0.58 (p=3.2E-10) | -0.32 (top 0.5%) |
| CAMK2A | -0.42 (p=1.1E-05) | -0.21 | -0.55 (p=4.0E-09) | -0.30 (top 0.5%) |
| TUBB2A | -0.54 (p=7.6E-09) | -0.46 (top 0.5%) | -0.48 (p=3.3E-07) | -0.28 (top 0.5%) |
| TUBA1A | -0.42 (p=1.4E-05) | -0.28 (top 0.5%) | -0.46 (p=1.4E-04) | -0.24 (top 1.0%) |

C: Correlations between Fluphenazine (VCP, 1uM-6h vs. 20 uM-24h) and Gene KD signatures.
| Gene KD | Low Dose, Short exposure:<br>LINCSCP_64588<br>(100) | Low Dose, Short exposure:<br>LINCSCP_64588<br>(978) | High Dose, Long exposure:<br>LINCSCP_64498<br>(100) | High Dose, Long exposure:<br>LINCSCP_64498<br>(978) |
| --- | --- | --- | --- | --- |
| VCAN | -0.67 (p=2.8E-14) | -0.40 (top 0.5%) | -0.35 (p=4.0E-04) | -0.34 (top 0.5%) |
| PRKCB | -0.54 (p=6.9E-09) | -0.36 (top 0.5%) | -0.41 (p=2.6E-06) | -0.33 (top 0.5%) |
| CAMK2A | -0.36 (p=2.0E-04) | -- | -0.27 (p=7.0E-03) | -- |
| TUBB2A | -0.62 (p=5.0E-12) | -0.43 (top 0.5%) | -0.33 (p=7.0E-04) | -0.31 (top 0.5%) |
| TUBA1A | -0.38 (p=1.0E-04) | -0.22 | -0.46 (p=1.3E-06) | -0.27 (top 0.5%) |

**Table S6.**
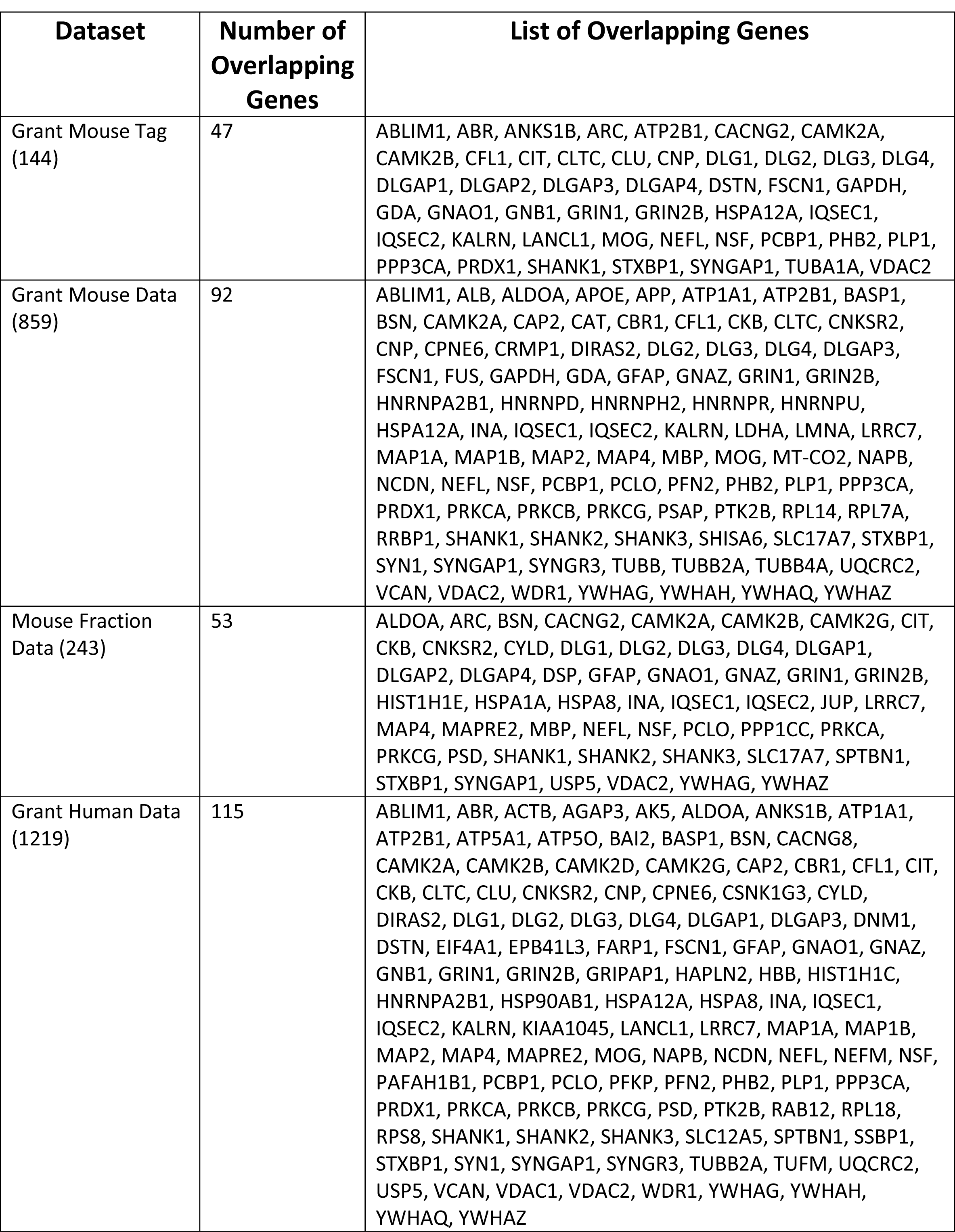
PSD-95 interactome overlap with literature derived datasets.

**Figure S1.**
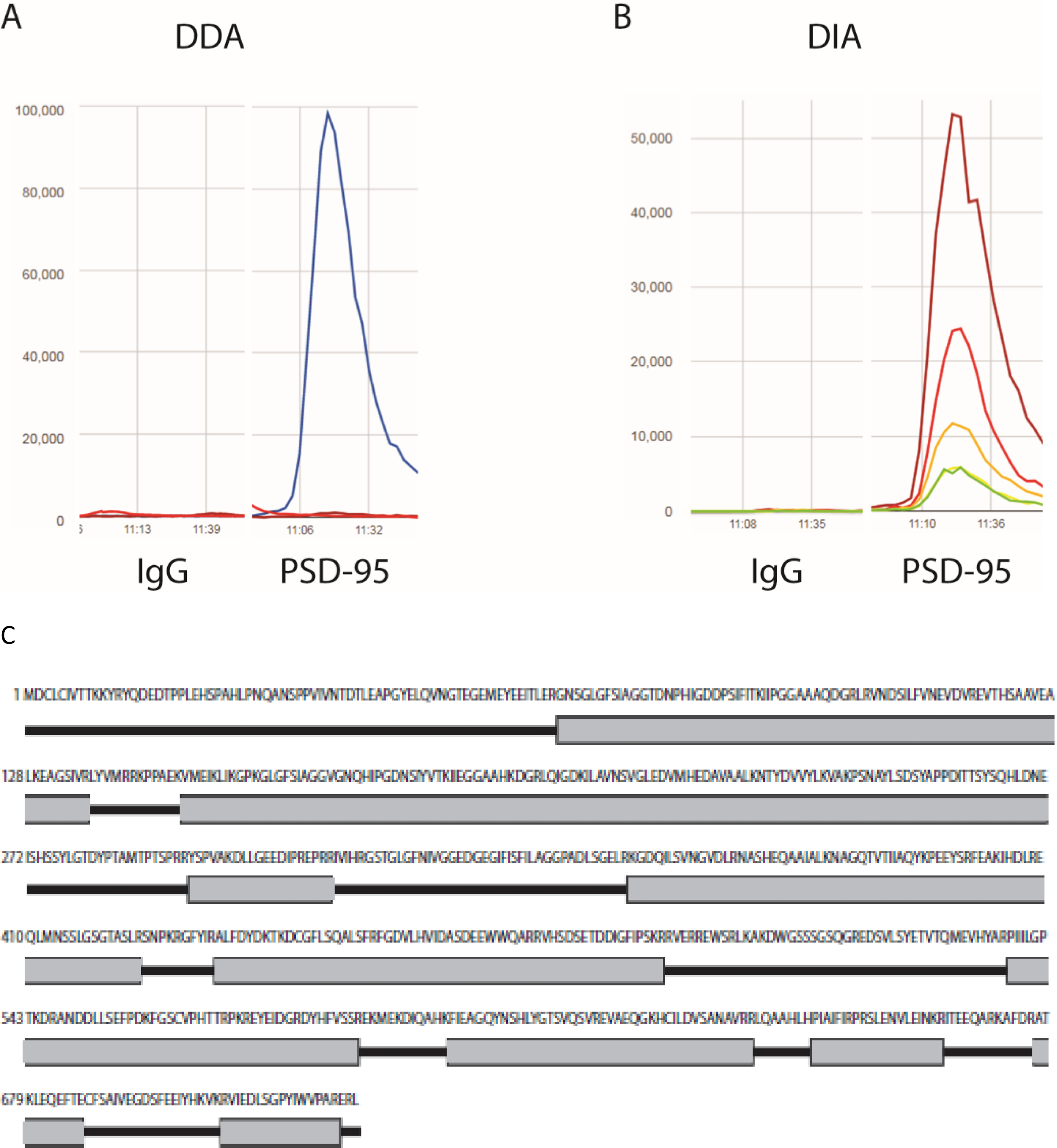
PSD-95 affinity purification specificity analysis. Data-dependent acquisition identification of a PSD-95 peptide in IgG vs PSD-95 isolation (A). Data-independent acquisition identification of a PSD-95 peptide in IgG vs PSD-95 isolation (B). Protein coverage for PSD-95 (C). 65.33% coverage (highlighted in grey) of human PSD-95 protein (Uniprot #P78352) was obtained from a single sample using a combination of data-dependent acquisition (DDA) and data independent acquisition (DIA) techniques on a liquid chromatography-mass spectrometry platform. Two replicates for each DDA and DIA were performed per single sample.

## Bibliography

1. Kandel ER, S.J., Jessell TM, Principles of Neural Science. 4th ed. 2000, New York: McGraw-Hill.

2. Zheng, W., et al., Neurocognitive dysfunction in subjects at clinical high risk for psychosis: A meta-analysis. J Psychiatr Res, 2018. 103: p. 38–45.

3. Gallo, F.T., et al., Immediate Early Genes, Memory and Psychiatric Disorders: Focus on c-Fos, Egr1 and Arc. Front Behav Neurosci, 2018. 12: p. 79.

4. Scheefhals, N. and H.D. MacGillavry, Functional organization of postsynaptic glutamate receptors. Mol Cell Neurosci, 2018.

5. Nanou, E. and W.A. Catterall, Calcium Channels, Synaptic Plasticity, and Neuropsychiatric Disease. Neuron, 2018. 98(3): p. 466–481.

6. Kim, S., H. Kim, and J.W. Um, Synapse development organized by neuronal activity-regulated immediate-early genes. Exp Mol Med, 2018. 50(4): p. 11.

7. Suh, Y.H., K. Chang, and K.W. Roche, Metabotropic glutamate receptor trafficking. Mol Cell Neurosci, 2018.

8. Bayes, A., et al., Characterization of the proteome, diseases and evolution of the human postsynaptic density. Nat Neurosci, 2011. 14(1): p. 19–21.

9. Chen, X., et al., Mass of the postsynaptic density and enumeration of three key molecules. Proc Natl Acad Sci U S A, 2005. 102(32): p. 11551–6.

10. Cheng, D., et al., Relative and absolute quantification of postsynaptic density proteome isolated from rat forebrain and cerebellum. Mol Cell Proteomics, 2006. 5(6): p. 1158–70.

11. Sugiyama, Y., et al., Determination of absolute protein numbers in single synapses by a GFP-based calibration technique. Nat Methods, 2005. 2(9): p. 677–84.

12. Albert, R., H. Jeong, and A.L. Barabasi, Error and attack tolerance of complex networks. Nature, 2000. 406(6794): p. 378–82.

13. Beique, J.C. and R. Andrade, PSD-95 regulates synaptic transmission and plasticity in rat cerebral cortex. J Physiol, 2003. 546(Pt 3): p. 859–67.

14. Ehrlich, I. and R. Malinow, Postsynaptic density 95 controls AMPA receptor incorporation during long-term potentiation and experience-driven synaptic plasticity. J Neurosci, 2004. 24(4): p. 916–27.

15. Elias, G.M., et al., Differential trafficking of AMPA and NMDA receptors by SAP102 and PSD-95 underlies synapse development. Proc Natl Acad Sci U S A, 2008. 105(52): p. 20953–8.

16. Schnell, E., et al., Direct interactions between PSD-95 and stargazin control synaptic AMPA receptor number. Proc Natl Acad Sci U S A, 2002. 99(21): p. 13902–7.

17. Stein, V., et al., Postsynaptic density-95 mimics and occludes hippocampal long-term potentiation and enhances long-term depression. J Neurosci, 2003. 23(13): p. 5503–6.

18. Ehrlich, I., et al., PSD-95 is required for activity-driven synapse stabilization. Proc Natl Acad Sci U S A, 2007. 104(10): p. 4176–81.

19. Elias, G.M., et al., Synapse-specific and developmentally regulated targeting of AMPA receptors by a family of MAGUK scaffolding proteins. Neuron, 2006. 52(2): p. 307–20.

20. Grant, S.G., Synaptopathies: diseases of the synaptome. Curr Opin Neurobiol, 2012. 22(3): p. 522–9.

21. Fromer, M., et al., De novo mutations in schizophrenia implicate synaptic networks. Nature, 2014. 506(7487): p. 179–84.

22. Coley, A.A. and W.J. Gao, PSD95: A synaptic protein implicated in schizophrenia or autism? Prog Neuropsychopharmacol Biol Psychiatry, 2018. 82: p. 187–194.

23. Bar-shira, O. and G. Chechik, Predicting protein-protein interactions in the post synaptic density. Mol Cell Neurosci, 2013. 56: p. 128–39.

24. Klemmer, P., A.B. Smit, and K.W. Li, Proteomics analysis of immuno-precipitated synaptic protein complexes. J Proteomics, 2009. 72(1): p. 82–90.

25. Emes, R.D. and S.G. Grant, The human postsynaptic density shares conserved elements with proteomes of unicellular eukaryotes and prokaryotes. Front Neurosci, 2011. 5: p. 44.

26. Callister, S.J., et al., Normalization approaches for removing systematic biases associated with mass spectrometry and label-free proteomics. J Proteome Res, 2006. 5(2): p. 277–86.

27. Valikangas, T., T. Suomi, and L.L. Elo, A systematic evaluation of normalization methods in quantitative label-free proteomics. Brief Bioinform, 2018. 19(1): p. 1–11.

28. Devarajan, K., Nonnegative matrix factorization: an analytical and interpretive tool in computational biology. PLoS Comput Biol, 2008. 4(7): p. e1000029.

29. Pascual-Montano, A., et al., bioNMF: a versatile tool for non-negative matrix factorization in biology. BMC Bioinformatics, 2006. 7: p. 366.

30. Wang, G., A.V. Kossenkov, and M.F. Ochs, LS-NMF: a modified non-negative matrix factorization algorithm utilizing uncertainty estimates. BMC Bioinformatics, 2006. 7: p. 175.

31. Heaven, M.R., et al., Systematic evaluation of data-independent acquisition for sensitive and reproducible proteomics-a prototype design for a single injection assay. J Mass Spectrom, 2016. 51(1): p. 1–11.

32. Adusumilli, R. and P. Mallick, Data Conversion with ProteoWizard msConvert. Methods Mol Biol, 2017. 1550: p. 339–368.

33. Holman, J.D., D.L. Tabb, and P. Mallick, Employing ProteoWizard to Convert Raw Mass Spectrometry Data. Curr Protoc Bioinformatics, 2014. 46: p. 13 24 1–9.

34. Kessner, D., et al., ProteoWizard: open source software for rapid proteomics tools development. Bioinformatics, 2008. 24(21): p. 2534–6.

35. Heaven, M.R., et al., Systematic evaluation of data-independent acquisition for sensitive and reproducible proteomics—a prototype design for a single injection assay. Journal of Mass Spectrometry, 2016. 51(1): p. 1–11.

36. Heaven, M.R., et al., Micro-Data-Independent Acquisition for High-Throughput Proteomics and Sensitive Peptide Mass Spectrum Identification. Anal Chem, 2018. 90(15): p. 8905–8911.

37. Heaven, M., et al., Systematic Evaluation of Data-Independent Acquisition for Sensitive and Reproducible Proteomics – a Prototype Design for a Single Injection Assay. J Mass Spectrom, 2015.

38. Vergeynst, L., H. Van Langenhove, and K. Demeestere, Balancing the false negative and positive rates in suspect screening with high-resolution Orbitrap mass spectrometry using multivariate statistics. Anal Chem, 2015. 87(4): p. 2170–7.

39. Kaimal, V., et al., ToppCluster: a multiple gene list feature analyzer for comparative enrichment clustering and network-based dissection of biological systems. Nucleic Acids Res, 2010. 38(Web Server issue): p. W96–102.

40. Shamsaei, B., et al., piNET: a versatile web platform for downstream analysis and visualization of proteomics data. bioRxiv, 2019.

41. Koleti, A., et al., Data Portal for the Library of Integrated Network-based Cellular Signatures (LINCS) program: integrated access to diverse large-scale cellular perturbation response data. Nucleic Acids Res, 2018. 46(D1): p. D558–D566.

42. Ong, E., et al., Ontological representation, integration, and analysis of LINCS cell line cells and their cellular responses. BMC Bioinformatics, 2017. 18(Suppl 17): p. 556.

43. Cheng, L. and L. Li, Systematic Quality Control Analysis of LINCS Data. CPT Pharmacometrics Syst Pharmacol, 2016. 5(11): p. 588–598.

44. Wang, Z., N.R. Clark, and A. Ma’ayan, Drug-induced adverse events prediction with the LINCS L1000 data. Bioinformatics, 2016. 32(15): p. 2338–45.

45. Liu, C., et al., Compound signature detection on LINCS L1000 big data. Mol Biosyst, 2015. 11(3): p. 714–22.

46. Duan, Q., et al., LINCS Canvas Browser: interactive web app to query, browse and interrogate LINCS L1000 gene expression signatures. Nucleic Acids Res, 2014. 42(Web Server issue): p. W449-60.

47. Vempati, U.D., et al., Metadata Standard and Data Exchange Specifications to Describe, Model, and Integrate Complex and Diverse High-Throughput Screening Data from the Library of Integrated Network-based Cellular Signatures (LINCS). J Biomol Screen, 2014. 19(5): p. 803–16.

48. Dosemeci, A., et al., Composition of the synaptic PSD-95 complex. Mol Cell Proteomics, 2007. 6(10): p. 1749–60.

49. Dunham, W.H., M. Mullin, and A.C. Gingras, Affinity-purification coupled to mass spectrometry: basic principles and strategies. Proteomics, 2012. 12(10): p. 1576–90.

50. Dunham, W.H., et al., A cost-benefit analysis of multidimensional fractionation of affinity purification-mass spectrometry samples. Proteomics, 2011. 11(13): p. 2603–12.

51. Morris, J.H., et al., Affinity purification-mass spectrometry and network analysis to understand protein-protein interactions. Nat Protoc, 2014. 9(11): p. 2539–54.

52. Varikuti, D.P., et al., Evaluation of non-negative matrix factorization of grey matter in age prediction. Neuroimage, 2018. 173: p. 394–410.

53. Greene, D., et al., Ensemble non-negative matrix factorization methods for clustering protein-protein interactions. Bioinformatics, 2008. 24(15): p. 1722–8.

54. Chalise, P. and B.L. Fridley, Integrative clustering of multi-level ‘omic data based on non-negative matrix factorization algorithm. PLoS One, 2017. 12(5): p. e0176278.

55. Benito, E., et al., The BET/BRD inhibitor JQ1 improves brain plasticity in WT and APP mice. Transl Psychiatry, 2017. 7(9): p. e1239.

56. Korb, E., et al., BET protein Brd4 activates transcription in neurons and BET inhibitor Jq1 blocks memory in mice. Nat Neurosci, 2015. 18(10): p. 1464–73.

57. McBrayer, Z.L., et al., Forebrain-Specific Loss of BMPRII in Mice Reduces Anxiety and Increases Object Exploration. PLoS One, 2015. 10(10): p. e0139860.

58. Osorio, C., et al., Growth differentiation factor 5 is a key physiological regulator of dendrite growth during development. Development, 2013. 140(23): p. 4751–62.

59. Zhang, D., et al., Development of bone morphogenetic protein receptors in the nervous system and possible roles in regulating trkC expression. J Neurosci, 1998. 18(9): p. 3314–26.

60. Soderstrom, S., H. Bengtsson, and T. Ebendal, Expression of serine/threonine kinase receptors including the bone morphogenetic factor type II receptor in the developing and adult rat brain. Cell Tissue Res, 1996. 286(2): p. 269–79.

61. De Wolf, H., et al., Transcriptional Characterization of Compounds: Lessons Learned from the Public LINCS Data. Assay Drug Dev Technol, 2016. 14(4): p. 252–60.

62. Zhang, W., et al., Structural basis of arc binding to synaptic proteins: implications for cognitive disease. Neuron, 2015. 86(2): p. 490–500.

63. Palomero-Gallagher, N., et al., Cytology and receptor architecture of human anterior cingulate cortex. J Comp Neurol, 2008. 508(6): p. 906–26.

64. Inoue, A. and S. Okabe, The dynamic organization of postsynaptic proteins: translocating molecules regulate synaptic function. Curr Opin Neurobiol, 2003. 13(3): p. 332–40.

65. Carlin, R.K., et al., Isolation and characterization of postsynaptic densities from various brain regions: enrichment of different types of postsynaptic densities. J Cell Biol, 1980. 86(3): p. 831–45.

66. Kelly, P.T. and C.W. Cotman, Synaptic proteins. Characterization of tubulin and actin and identification of a distinct postsynaptic density polypeptide. J Cell Biol, 1978. 79(1): p. 173–83.

67. Blomberg, F., R.S. Cohen, and P. Siekevitz, The structure of postsynaptic densities isolated from dog cerebral cortex. II. Characterization and arrangement of some of the major proteins within the structure. J Cell Biol, 1977. 74(1): p. 204–25.

68. Cohen, R.S., et al., The structure of postsynaptic densities isolated from dog cerebral cortex. I. Overall morphology and protein composition. J Cell Biol, 1977. 74(1): p. 181–203.

69. Banker, G., L. Churchill, and C.W. Cotman, Proteins of the postsynaptic density. J Cell Biol, 1974. 63(2 Pt 1): p. 456–65.

70. Cotman, C.W., et al., Isolation of postsynaptic densities from rat brain. J Cell Biol, 1974. 63(2 Pt 1): p. 441–55.

71. Cotman, C.W. and D. Taylor, Isolation and structural studies on synaptic complexes from rat brain. J Cell Biol, 1972. 55(3): p. 696–711.

72. Fernandez, E., et al., Targeted tandem affinity purification of PSD-95 recovers core postsynaptic complexes and schizophrenia susceptibility proteins. Mol Syst Biol, 2009. 5: p. 269.

73. Dosemeci, A., et al., Preparation of postsynaptic density fraction from hippocampal slices and proteomic analysis. Biochem Biophys Res Commun, 2006. 339(2): p. 687–94.

74. Vinade, L., et al., Affinity purification of PSD-95-containing postsynaptic complexes. J Neurochem, 2003. 87(5): p. 1255–61.

75. Zhu, F., et al., Architecture of the Mouse Brain Synaptome. Neuron, 2018. 99(4): p. 781–799 e10.

76. Roy, M., et al., Regional Diversity in the Postsynaptic Proteome of the Mouse Brain. Proteomes, 2018. 6(3).

77. Roy, M., et al., Proteomic analysis of postsynaptic proteins in regions of the human neocortex. Nat Neurosci, 2018. 21(1): p. 130–138.

78. Yuan, D., et al., Nest-building activity as a reproducible and long-term stroke deficit test in a mouse model of stroke. Brain Behav, 2018. 8(6): p. e00993.

79. Duszczyk, M., et al., Behavioral evaluation of ischemic damage to CA1 hippocampal neurons: effects of preconditioning. Acta Neurobiol Exp (Wars), 2006. 66(4): p. 311–9.

80. Kim, C., Nest building, general activity, and salt preference of rats following hippocampal ablation. J Comp Physiol Psychol, 1960. 53: p. 11–6.

81. Rubin, R.D., et al., Dynamic Hippocampal and Prefrontal Contributions to Memory Processes and Representations Blur the Boundaries of Traditional Cognitive Domains. Brain Sci, 2017. 7(7).

82. Rubin, R.D., et al., The role of the hippocampus in flexible cognition and social behavior. Front Hum Neurosci, 2014. 8: p. 742.

83. Duff, M.C., et al., Hippocampal amnesia disrupts creative thinking. Hippocampus, 2013. 23(12): p. 1143–9.

84. Rubin, R.D., et al., How do I remember that I know you know that I know? Psychol Sci, 2011. 22(12): p. 1574–82.

85. Klaassen, R.V., et al., Shisa6 traps AMPA receptors at postsynaptic sites and prevents their desensitization during synaptic activity. Nat Commun, 2016. 7: p. 10682.

86. Tokue, M., et al., SHISA6 Confers Resistance to Differentiation-Promoting Wnt/beta-Catenin Signaling in Mouse Spermatogenic Stem Cells. Stem Cell Reports, 2017. 8(3): p. 561–575.

87. Abeliovich, A., et al., PKC gamma mutant mice exhibit mild deficits in spatial and contextual learning. Cell, 1993. 75(7): p. 1263–71.

88. Abeliovich, A., et al., Modified hippocampal long-term potentiation in PKC gamma-mutant mice. Cell, 1993. 75(7): p. 1253–62.

89. Patten, S.A., et al., Protein kinase Cgamma is a signaling molecule required for the developmental speeding of alpha-amino-3-hydroxyl-5-methyl-4-isoxazole-propionate receptor kinetics. Eur J Neurosci, 2010. 31(9): p. 1561–73.

90. Patten, S.A. and D.W. Ali, PKCgamma-induced trafficking of AMPA receptors in embryonic zebrafish depends on NSF and PICK1. Proc Natl Acad Sci U S A, 2009. 106(16): p. 6796–801.

91. Patten, S.A., et al., Differential expression of PKC isoforms in developing zebrafish. Int J Dev Neurosci, 2007. 25(3): p. 155–64.

92. Hanley, J.G., et al., NSF ATPase and alpha-/beta-SNAPs disassemble the AMPA receptor-PICK1 complex. Neuron, 2002. 34(1): p. 53–67.

93. Araki, Y., D.T. Lin, and R.L. Huganir, Plasma membrane insertion of the AMPA receptor GluA2 subunit is regulated by NSF binding and Q/R editing of the ion pore. Proc Natl Acad Sci U S A, 2010. 107(24): p. 11080–5.

94. Lin, D. and R. Huganir, PICK1 and phosphorylation of the glutamate receptor 2 (GluR2) AMPA receptor subunit regulates GluR2 recycling after NMDA receptor-induced internalization. J Neurosci, 2007. 27(50): p. 13903–8.

95. Huang, Y., et al., S-nitrosylation of N-ethylmaleimide sensitive factor mediates surface expression of AMPA receptors. Neuron, 2005. 46(4): p. 533–40.

96. Gardner, S.M., et al., Calcium-permeable AMPA receptor plasticity is mediated by subunit-specific interactions with PICK1 and NSF. Neuron, 2005. 45(6): p. 903–15.

97. Steinberg, J.P., R.L. Huganir, and D.J. Linden, N-ethylmaleimide-sensitive factor is required for the synaptic incorporation and removal of AMPA receptors during cerebellar long-term depression. Proc Natl Acad Sci U S A, 2004. 101(52): p. 18212–6.

98. Song, I., et al., Interaction of the N-ethylmaleimide-sensitive factor with AMPA receptors. Neuron, 1998. 21(2): p. 393–400.

99. Tewari, R., et al., Armadillo-repeat protein functions: questions for little creatures. Trends Cell Biol, 2010. 20(8): p. 470–81.

100. D’Souza, C.A., et al., Identification of a set of genes showing regionally enriched expression in the mouse brain. BMC Neurosci, 2008. 9: p. 66.

101. Walikonis, R.S., et al., Densin-180 forms a ternary complex with the (alpha)-subunit of Ca2+/calmodulin-dependent protein kinase II and (alpha)-actinin. J Neurosci, 2001. 21(2): p. 423–33.

102. Strack, S., et al., Association of calcium/calmodulin-dependent kinase II with developmentally regulated splice variants of the postsynaptic density protein densin-180. J Biol Chem, 2000. 275(33): p. 25061–4.

103. Kennedy, M.B., Signal transduction molecules at the glutamatergic postsynaptic membrane. Brain Res Brain Res Rev, 1998. 26(2-3): p. 243–57.

104. Kennedy, M.B., The postsynaptic density at glutamatergic synapses. Trends Neurosci, 1997. 20(6): p. 264–8.

105. Apperson, M.L., I.S. Moon, and M.B. Kennedy, Characterization of densin-180, a new brain-specific synaptic protein of the O-sialoglycoprotein family. J Neurosci, 1996. 16(21): p. 6839–52.

106. Ohtakara, K., et al., Densin-180, a synaptic protein, links to PSD-95 through its direct interaction with MAGUIN-1. Genes Cells, 2002. 7(11): p. 1149–60.

107. Vessey, J.P. and D. Karra, More than just synaptic building blocks: scaffolding proteins of the post-synaptic density regulate dendritic patterning. J Neurochem, 2007. 102(2): p. 324–32.

108. Quitsch, A., et al., Postsynaptic shank antagonizes dendrite branching induced by the leucine-rich repeat protein Densin-180. J Neurosci, 2005. 25(2): p. 479–87.

109. Kim, K., et al., Dendrite-like process formation and cytoskeletal remodeling regulated by delta-catenin expression. Exp Cell Res, 2002. 275(2): p. 171–84.

110. Jones, S.B., et al., Glutamate-induced delta-catenin redistribution and dissociation from postsynaptic receptor complexes. Neuroscience, 2002. 115(4): p. 1009–21.

111. Carlisle, H.J., et al., Deletion of densin-180 results in abnormal behaviors associated with mental illness and reduces mGluR5 and DISC1 in the postsynaptic density fraction. J Neurosci, 2011. 31(45): p. 16194–207.

112. Sardina, J.M., et al., Amelioration of the typical cognitive phenotype in a patient with the 5pter deletion associated with Cri-du-chat syndrome in addition to a partial duplication of CTNND2. Am J Med Genet A, 2014. 164A(7): p. 1761–4.

113. Arikkath, J., et al., Delta-catenin regulates spine and synapse morphogenesis and function in hippocampal neurons during development. J Neurosci, 2009. 29(17): p. 5435–42.

114. Israely, I., et al., Deletion of the neuron-specific protein delta-catenin leads to severe cognitive and synaptic dysfunction. Curr Biol, 2004. 14(18): p. 1657–63.

115. Medina, M., et al., Hemizygosity of delta-catenin (CTNND2) is associated with severe mental retardation in cri-du-chat syndrome. Genomics, 2000. 63(2): p. 157–64.

116. Gilbert, J. and H.Y. Man, The X-Linked Autism Protein KIAA2022/KIDLIA Regulates Neurite Outgrowth via N-Cadherin and delta-Catenin Signaling. eNeuro, 2016. 3(5).

117. Turner, T.N., et al., Loss of delta-catenin function in severe autism. Nature, 2015. 520(7545): p. 51–6.

118. Brigidi, G.S., et al., Activity-regulated trafficking of the palmitoyl-acyl transferase DHHC5. Nat Commun, 2015. 6: p. 8200.

119. Brigidi, G.S., et al., Palmitoylation of delta-catenin by DHHC5 mediates activity-induced synapse plasticity. Nat Neurosci, 2014. 17(4): p. 522–32.

120. Blitzer, R.D., R. Iyengar, and E.M. Landau, Postsynaptic signaling networks: cellular cogwheels underlying long-term plasticity. Biol Psychiatry, 2005. 57(2): p. 113–9.

121. Kristiansen, L.V., et al., NMDA receptors and schizophrenia. Curr Opin Pharmacol, 2007. 7(1): p. 48–55.

122. Gardoni, F., E. Marcello, and M. Di Luca, Postsynaptic density-membrane associated guanylate kinase proteins (PSD-MAGUKs) and their role in CNS disorders. Neuroscience, 2009. 158(1): p. 324–33.

123. Grant, S.G., The molecular evolution of the vertebrate behavioural repertoire. Philos Trans R Soc Lond B Biol Sci, 2016. 371(1685): p. 20150051.

124. Kristiansen, L.V., et al., Changes in NMDA receptor subunits and interacting PSD proteins in dorsolateral prefrontal and anterior cingulate cortex indicate abnormal regional expression in schizophrenia. Mol Psychiatry, 2006. 11(8): p. 737-47, 705.

125. Kristiansen, L.V. and J.H. Meador-Woodruff, Abnormal striatal expression of transcripts encoding NMDA interacting PSD proteins in schizophrenia, bipolar disorder and major depression. Schizophr Res, 2005. 78(1): p. 87–93.

126. Li, J.M., et al., Role of the DLGAP2 gene encoding the SAP90/PSD-95-associated protein 2 in schizophrenia. PLoS One, 2014. 9(1): p. e85373.

127. Rasmussen, A.H., H.B. Rasmussen, and A. Silahtaroglu, The DLGAP family: neuronal expression, function and role in brain disorders. Mol Brain, 2017. 10(1): p. 43.

128. Devinsky, O., M.J. Morrell, and B.A. Vogt, Contributions of anterior cingulate cortex to behaviour. Brain, 1995. 118 (Pt 1): p. 279–306.

129. Allman, J.M., et al., The anterior cingulate cortex. The evolution of an interface between emotion and cognition. Ann N Y Acad Sci, 2001. 935: p. 107–17.

130. Egles, C., et al., Laminins containing the beta2 chain modulate the precise organization of CNS synapses. Mol Cell Neurosci, 2007. 34(3): p. 288–98.

131. Bayes, A., et al., Comparative study of human and mouse postsynaptic proteomes finds high compositional conservation and abundance differences for key synaptic proteins. PLoS One, 2012. 7(10): p. e46683.

132. Radner, S., et al., beta2 and gamma3 laminins are critical cortical basement membrane components: ablation of Lamb2 and Lamc3 genes disrupts cortical lamination and produces dysplasia. Dev Neurobiol, 2013. 73(3): p. 209–29.

133. Sharma, T. and M.I. Siddiqi, In silico identification and design of potent peptide inhibitors against PDZ-3 domain of Postsynaptic Density Protein (PSD-95). J Biomol Struct Dyn, 2018: p. 1–13.

134. Furuyashiki, T., et al., Citron, a Rho-target, interacts with PSD-95/SAP-90 at glutamatergic synapses in the thalamus. J Neurosci, 1999. 19(1): p. 109–18.

135. Zhang, W., et al., Citron binds to PSD-95 at glutamatergic synapses on inhibitory neurons in the hippocampus. J Neurosci, 1999. 19(1): p. 96–108.

136. Johnstone, M., et al., Copy Number Variations in DISC1 and DISC1-Interacting Partners in Major Mental Illness. Mol Neuropsychiatry, 2015. 1(3): p. 175–190.

137. Kirov, G., et al., The penetrance of copy number variations for schizophrenia and developmental delay. Biol Psychiatry, 2014. 75(5): p. 378–85.

138. Grozeva, D., et al., Reduced burden of very large and rare CNVs in bipolar affective disorder. Bipolar Disord, 2013. 15(8): p. 893–8.

139. Kirov, G., et al., De novo CNV analysis implicates specific abnormalities of postsynaptic signalling complexes in the pathogenesis of schizophrenia. Mol Psychiatry, 2012. 17(2): p. 142–53.

140. Feinberg, A.P. and S.H. Snyder, Phenothiazine drugs: structure-activity relationships explained by a conformation that mimics dopamine. Proc Natl Acad Sci U S A, 1975. 72(5): p. 1899–903.

141. Gu, W.H., et al., Requirement of PSD-95 for dopamine D1 receptor modulating glutamate NR1a/NR2B receptor function. Acta Pharmacol Sin, 2007. 28(6): p. 756–62.

142. Zhang, J., et al., Inhibition of the dopamine D1 receptor signaling by PSD-95. J Biol Chem, 2007. 282(21): p. 15778–89.

143. Sun, P., et al., PSD-95 regulates D1 dopamine receptor resensitization, but not receptor-mediated Gs-protein activation. Cell Res, 2009. 19(5): p. 612–24.

144. Zhang, J., et al., PSD-95 uncouples dopamine-glutamate interaction in the D1/PSD-95/NMDA receptor complex. J Neurosci, 2009. 29(9): p. 2948–60.

145. Cohen-Armon, M., A PARP1-Erk2 synergism is required for stimulation-induced expression of immediate early genes. Gene Transl Bioinform, 2016. 2.

146. Visochek, L., et al., A PARP1-ERK2 synergism is required for the induction of LTP. Sci Rep, 2016. 6: p. 24950.

